# MLL4 is required for the first embryonic collective cell migration whereas MLL3 is not required until birth

**DOI:** 10.1101/870329

**Authors:** Deepthi Ashokkumar, Qinyu Zhang, Christian Much, Anita S. Bledau, Jun Fu, Konstantinos Anastassiadis, A. Francis Stewart, Andrea Kranz

## Abstract

Methylation of histone 3 lysine 4 (H3K4) is a major epigenetic system associated with gene expression. In mammals there are six H3K4 methyltransferases related to yeast Set1 and fly Trithorax, including two orthologs of fly Trithorax-related: MLL3 and MLL4. Exome sequencing has documented high frequencies of *Mll3* and *Mll4* mutations in many types of human cancer. Despite this emerging importance, the requirements of these sister genes in mammalian development have only been incompletely reported. Here we examined the null phenotypes to establish that MLL3 is first required for lung maturation whereas MLL4 is first required for migration of the anterior visceral endoderm (AVE) that initiates gastrulation and is the first collective cell migration in development. This migration is preceded by a columnar to squamous transition in visceral endoderm cells that depends on MLL4. Furthermore, *Mll4* mutants display incompletely penetrant, sex distorted, embryonic haploinsufficiency and adult heterozygous mutants show aspects of Kabuki syndrome, indicating that MLL4 action, unlike MLL3, is dosage dependent. The highly specific and discordant functions of these sister genes argues against their action as general enhancer factors.

**Summary statement:** The H3K4 methyltransferases MLL3 and MLL4 have strikingly different null phenotypes during mouse development; MLL3 is required for lung maturation whereas MLL4 is required for anterior visceral endoderm migration.

## Introduction

The lysine methylation status of the histone 3 tail is central to epigenetic regulation pivoting on methylation of lysines at positions 4, 9, 27 and 36. All active Pol II promoters are characterized by trimethylation of histone 3 lysine 4 (H3K4me3) on the first nucleosome in the transcribed region. Dimethylation (H3K4me2) is a general characteristic of transcribed regions whereas monomethylation (H3K4me1) is a general characteristic of active chromatin with peaks on enhancers (Bannister and Kouzarides, 2011).

Mammals have six Set1/Trithorax type H3K4 methyltransferases in three pairs; SETD1A and B (KMT2F and KMT2G), which are homologues of yeast Set1; MLL1 and 2 (KMT2A and KMT2B), which are homologues of *Drosophila* Trithorax; and MLL3 and 4 (KMT2C and KMT2D), which are homologues of *Drosophila* Lost PHD fingers of Trr (*Ltr*) fused to Trithorax-related (*Trr*). All six are found in individual complexes however all six complexes share the same highly conserved scaffold, first reported for yeast Set1C (Roguev et al., 2001), composed of four subunits, WDR5, RBBP5, ASH2L, DPY30 termed ‘WRAD’ (Lee et al., 2006; Ruthenburg et al., 2007; Ernst and Vakoc, 2012), which surrounds the SET domain and is required for enzymatic activity (Kim et al., 2013; Hsu et al., 2018; Qu et al., 2018). SETD1A apparently conveys most H3K4me3 in most mammalian cell types (Bledau et al., 2014). Similarly the dSet1 homologue conveys most H3K4me3 in most *Drosophila* cell types (Mohan et al., 2011; Ardehali et al., 2011; Hallson et al., 2012). Consequently the Set1 homologues are primarily implicated in trimethylation and general promoter function. In contrast, evidence indicating that MLL3 and 4 are monomethylases (Weirich et al., 2015; Zhang et al., 2015; Li et al., 2016) has triggered their linkage to enhancer function (Lee et al., 2013; Rao and Dou, 2015; Piunti and Shilatifard, 2016). Whether they proceed to catalyze di- and tri-H3K4 methylation remains uncertain (Dhar et al., 2012) and the emergent model relating Set1 activities to promoters and MLL3/4 to enhancers requires further substantiation.

Histone 3 lysine methyltransferases are prominent members of both Trithorax-(Trx-G) and Polycomb-Groups (Pc-G) (Steffen and Ringrose, 2014; Schuettengruber et al., 2017) with the genetic opposition between Trx-G and Pc-G being exerted, in part, by a competition for the methylation status of the histone 3 tail on key nucleosomes (Schmitges et al., 2011). This opposition is central to epigenetic regulation in development, differentiation, homeostasis and more recently, oncogenesis (Chi et al., 2010; Rao and Dou, 2015; Soshnev et al., 2016) with several Trx-G and Pc-G factors, including the H3K27 methyltransferase, EZH2, implicated as oncogenes or tumor suppressors in a variety of malignancies. *Mll1* was discovered as the major leukemia gene at the 11q23.1 translocation involved in early onset childhood leukemia (Li and Ernst, 2014). The N-terminal half of MLL1 fused to many (now more than 70) different C-terminal partners, including AF4 and AF9 (Slany, 2009; Meyer et al., 2018) is leukemiogenic without the need for secondary mutations (Dobson et al., 2000). These MLL1 fusion proteins promote both acute lymphocytic (ALL) and acute myeloid (AML) leukemias, collectively termed mixed lineage leukemias.

Massively parallel sequencing of cancer exomes by the international cancer genome projects revealed somatic mutations in *Mll3* and *Mll4* in almost all cancers analyzed (Rao and Dou, 2015). Inactivating heterozygous mutations have been identified in patients with medulloblastoma, B cell lymphoma, bladder carcinoma, renal carcinoma and colorectal cancer, amongst many other cancers (Morin et al., 2011; Parsons et al., 2011; Pasqualucci et al., 2011). An explanation of these findings is lacking however recent evidence suggests that mutation of *Mll4* promotes defective transcription-coupled DNA repair (Kantidakis et al., 2016).

Exome sequencing also revealed mutations in *Mll4* as the cause of Kabuki syndrome type I (Ng et al., 2010; Li et al., 2011). All *Mll4* Kabuki mutations are apparently *de novo* somatic heterozygous nonsense or frameshift mutations that appear throughout the gene, but most commonly in exon 48. Most of these *Mll4* mutations truncate the protein and all are haploinsufficient (Banka et al., 2012; Bogershausen et al., 2015; Faundes et al., 2019). The less common Kabuki syndrome type 2 is caused by mutations of *Utx*. UTX, which is an H3K27 demethylase, is a subunit of the MLL4 complex (Lee et al., 2006; Lederer et al., 2012; Lederer et al., 2014; Banka et al., 2015). As for MLL1 and MLL2 (Denissov et al., 2014), MLL3 and MLL4 may have overlapping and redundant functions in mammalian cells (Lee et al., 2013). Notably, the H3K4 methyltransferase activities of MLL3 and MLL4 are dispensable for gene expression in mouse embryonic stem cells (ESCs) (Dorighi et al., 2017). Similarly, the catalytic activity of the SET domain of Trr (the fly homologue of MLL3/MLL4) is dispensable for development and viability in *Drosophila* (Rickels et al., 2017).

Unlike the other four H3K4 methyltransferases (Yagi et al., 1998; Ernst et al., 2004; Glaser et al., 2006; Jude et al., 2007; Glaser et al., 2009; Andreu-Vieyra et al., 2010; Bledau et al., 2014) and despite their emerging importance for cancer, the roles of *Mll3* and *Mll4* in mammalian development have only been partly described (Ang et al., 2016; Lee et al., 2013; Jang et al., 2017). Here we compare the null phenotypes of these two genes in mouse development.

## Material and Methods

### Targeting constructs

The targeting constructs for *Mll3* and *Mll4* were generated using Red/ET recombineering (Fu et al., 2010). For *Mll3* an FRT-SA-GT0-T2A-lacZneo-CoTC-FRT-loxP cassette was inserted into intron 48. Additionally, a loxP-rox-PGK-Blasticidin-pA-rox cassette was introduced into intron 49. For *Mll4*, a loxP site was introduced into intron 4 using a loxP-zeo-loxP cassette with subsequent removal in *E. coli* by Cre recombination using pSC101-BAD-Cre-tet (Anastassiadis et al., 2009). Then a loxP-FRT-SA-IRES-lacZneo-pA-FRT cassette was inserted in intron 1 of the gene. The homology arms were 5’ 4.6kb/3’ 5kb and 5’ 4.7kb/3’ 4.9kb for the *Mll3* and *Mll4* targeting constructs, respectively.

### Gene targeting and generation of conditional knockout mice

R1 ESCs were cultured with FCS-based medium (DMEM + GlutaMAX^TM^ (Invitrogen), 15% FCS (Fetal Calf Serum, PAA), 2 mM L-Glutamine (Invitrogen), 1xnon-essential amino acids (Invitrogen), 1 mM sodium pyruvate (Invitrogen), 0.1 mM β-mercaptoethanol, in the presence of LIF)) on Mitomycin-C inactivated mouse embryonic fibroblasts. Cells (1x10^7^) were electroporated with 40 μg of the linearized targeting construct using standard conditions (250 V, 500 μF) and selected with 0.2 mg/ml G418. Correct integration in the *Mll3* locus was confirmed by Southern blot analysis using 5’ and 3’ external probes (4 positive clones out of 36). Correct integration in the *Mll4* locus was confirmed by Southern blot analysis using an internal probe and 5’ and 3’ external probes (6 positive clones out of 32). To remove the additional selection cassette (PGK-Blasticidin-polyA) downstream of exon 49, correctly targeted *Mll3* clones were electroporated with CAGGS-Dre-IRES-puro expression vector (Anastassiadis et al., 2009) and clones screened by PCR for complete recombination and sensitivity to blasticidin. For both *Mll3* and *Mll4* two correctly targeted ES cell clones were injected into blastocysts. For *Mll3* only one of the clones gave germ line transmission. *Mll3^D/+^* mice were crossed to the *CAGGs-Flpo* line (Kranz et al., 2010) to generate *Mll3^FD/+^* mice. Subsequently, those mice were crossed to the *PGK-Cre* line (Lallemand et al., 1998) to produce *Mll3^FDC/+^* mice. *Mll3^D/+^* and *Mll3^FDC/+^* mice were backcrossed at least 15 generations to C57BL/6JOlaHsd mice. Regarding *Mll4* both clones gave rise to several chimeras but only one of these males was able to establish germ line transmission. *Mll4^A/+^* mice were backcrossed at least 15 generations to CD1 mice. Primers for genotyping are provided in Table S1. All animal experiments were performed according to German law.

### Western blot analysis

ESCs were homogenized in buffer E (20 mM HEPES pH 8.0, 350 mM NaCl, 10% glycerol, 0.1% Tween 20, 1 mM PMSF, 1x complete protease inhibitor cocktail) and protein extracts were obtained after three cycles of freezing and thawing. Whole cell extracts were subsequently separated by NuPAGE 3-8% Tris-acetate gel (Invitrogen), transferred to PVDF membranes and probed with an MLL4 specific polyclonal antibody. The antibody was generated by immunizing rabbits with a mixture of three KLH-conjugated synthetic peptides from the central part of the MLL4 protein (QRPRFYPVSEELHRLAP, NGDEFDLLAYT, KQQLSAQTQRLAPS) (extended data file 1). Antibody information and dilution can be found in Table S3.

### Whole-mount X-gal staining and immunostaining

Embryos were dissected, fixed with 0.2% glutaraldehyde and X-gal staining was performed as described (Kranz et al., 2010). Embryos and organs were dissected and fixed with 4% paraformaldehyde overnight. Dehydration and paraffin infiltration was done automatically using the Paraffin-Infiltration-Processor (STP 420, Zeiss). The dehydrated tissues were embedded in paraffin (Paraffin Embedding Center EG116, Leica) and sections were prepared. Antigen retrieval was performed by microwaving slides in 10 mM citrate buffer (pH 6.0) for 12 min (Microwave RHS 30, Diapath). Immunohistochemical staining was carried out as previously described (Bledau et al., 2014). DAB images were collected with an Olympus WF upright microscope. For immunofluorescence procedures, the sections were permeabilized in 0.5% Triton X-100 in PBS for 10 min, blocked for 1 hour at RT, incubated with primary antibody overnight at 4°C, followed by secondary antibody for 2 hours at RT. Sections were mounted with Mowiol and imaged with Zeiss Scanning confocal microscope LSM/780. Antibody information and dilutions can be found in Table S3.

### Whole-mount in situ hybridization

Whole-mount *in situ* hybridization was carried out according to standard procedures (Riddle et al., 1993; Piette et al., 2008). Digoxigenin-labelled riboprobes specific for the following genes were used in this study: *Brachyury* (Herrmann, 1991), *Mox1*, *Hoxb1 and Wnt1* (Glaser et al., 2006), *Otx2* (Ang et al., 1996)*, Nodal, Eomes, Bmp4 and Hex* (Norris et al., 2002), *Cer1 (Belo et al., 1997), Foxa2 (Norris et al., 2002)* and *Gsc, Lhx1* and *Wnt3* (Liu et al., 1999), *Dkk1* (Stuckey et al., 2011), *Lefty1* (Stuckey et al., 2011), *Shh* and *Tbx6* (Alten et al., 2012). Anti-dig-AP antibody and NBT/BCIP colorimetric signal detection were used for whole-mount *in situ* hybridizations. Embryos were imaged with Nikon SMZ 1500 stereomicroscope.

### Whole-mount immunofluorescence for Hex

For whole-mount immunofluorescence, PFA-fixed embryos were permeabilized in 0.5% Triton X-100 in PBS for 1 hour at RT, incubated with anti-Hex antibody (Hoshino et al., 2015) overnight at 4°C followed by goat anti-rabbit IgG-CFL 488 (Santa Cruz) secondary antibody. The embryos were imaged with Zeiss Scanning confocal microscope LSM/780.

### Glucose and insulin tolerance test

For glucose tolerance test, mice fasted 16 h before 1.5 mg glucose per g body weight was applied by gavage. For insulin tolerance test, mice were injected intraperitoneally with 0.75 mU insulin per g body weight after 6 h of fasting. For measurement of blood glucose levels blood samples were drawn at 0, 15, 30, 60, 90, and 120 min after administration of the respective solution.

### Reverse transcription and quantitative PCR (qRT-PCR) analysis

Total RNA was isolated using Trizol reagent (Sigma-Aldrich) and reverse transcribed using the AffinityScript Multiple Temperature cDNA Synthesis kit (Agilent Technologies). Real-time quantitative PCR was performed with Go Taq qPCR Master Mix (Promega) by Mx3000P QPCR System (Agilent Technologies). Ct values were normalized against *Rpl19*. Primer sequences and length of the amplified products are given in Table S2. Fold differences in expression levels were calculated according to the 2^−ΔCt^ method (Livak and Schmittgen, 2001).

## Results

### Mll3 and Mll4 are conserved paralogs

MLL3 and MLL4 are the largest known nuclear proteins at 4903 and 5588 amino acids respectively and their genes clearly arose by duplication. Both genes and proteins have the same architecture based on the positions of splice sites, PHD fingers, HMG box, FYRN/FYRC and SET domains (Fig. 1A; extended data file 1). Except for PHD3 of MLL3, which has been lost from MLL4 (because PHD3 can be found in *Drosophila Lpt* (Lost PHD fingers of *Trr*)) (Mohan et al., 2011; Chauhan et al., 2012), the other five PHD and ePHD fingers share high identity. Both proteins are notable for their extensive regions of low sequence complexity, particularly MLL4 contains several lengthy stretches of glutamine repeats including a patch of 450 amino acids C-terminal to the HMG box with more than 50% glutamines and a 600 amino acid patch with one-third prolines after its second PHD finger (extended data file 1). Both genes encompass more than 50 exons (Fig. 1B, Fig. 1C) spliced to very long mRNAs; 14 kb for *Mll3* and 19 kb for *Mll4,* that are widely expressed in the embryo (Fig. S1C, S2C).

**Fig. 1.**
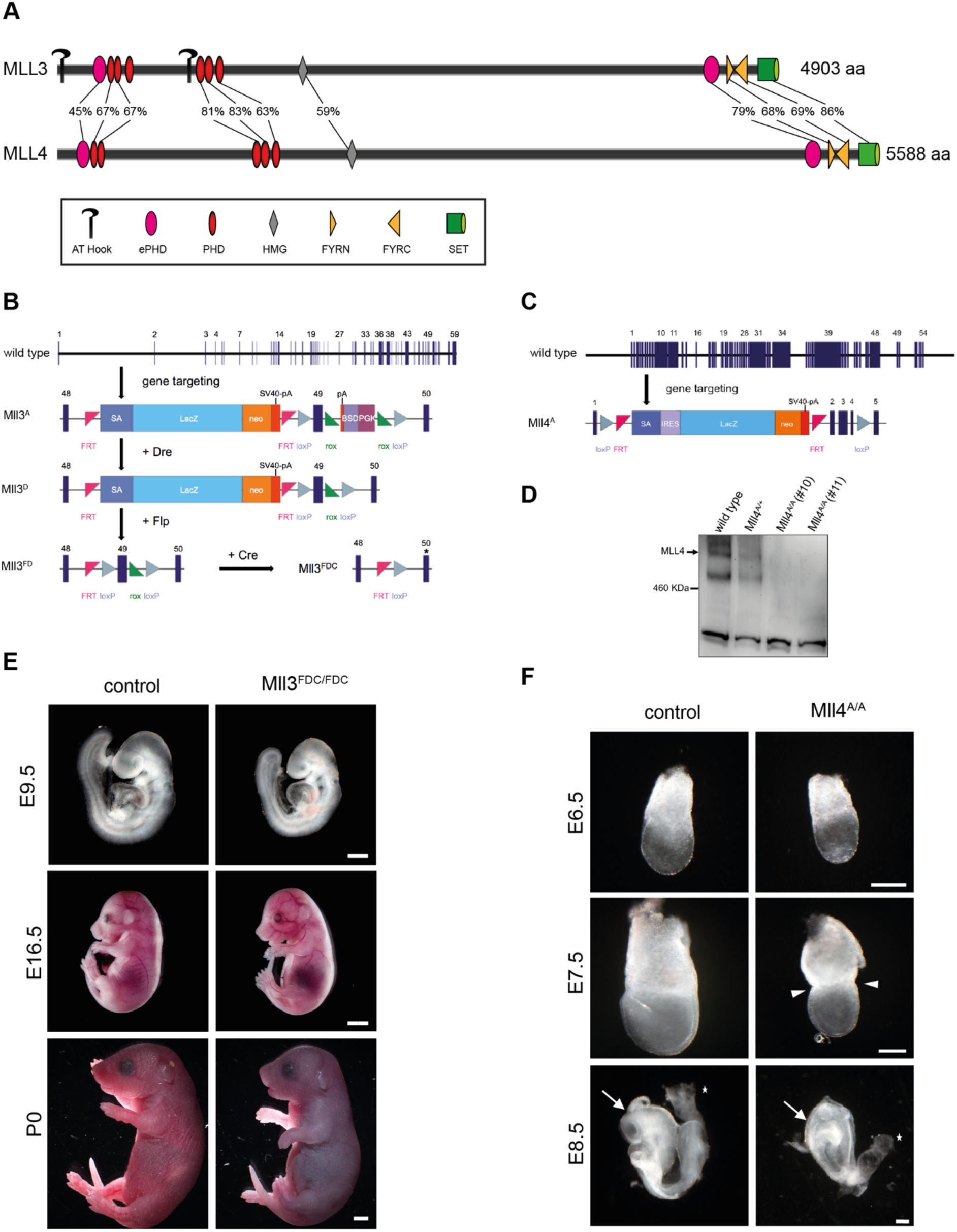
Allele design and embryonic phenotype of the *Mll3 and Mll4* mutant embryos. (A) Amino acid sequence alignment of murine MLL3 and MLL4 based on EMBOSS Needle. FYRN and FYRC = F/Y-rich N- or C-terminus, HMG = high mobility group, ePHD = extended PHD finger. (B) Diagram of the *Mll3* knockout first allele (*Mll3^A^*). The *Mll3* wild type (wt) allele contains 59 coding exons. The targeted allele, *Mll3^A^*, is similar to the Dre recombined allele, *Mll3^D^*, with the PGK-Blasticidin selection cassette removed. The *Mll3^D^* allele is converted to conditional (*Mll3^FD^*) upon FLP recombination. Cre recombination leads to excision of the frameshifting exon 49 generating the conditional mutant allele (*Mll3^FDC^*). Blue rectangles with numbers on top indicate exons. The star depicts the exon with the premature stop codon. SA = splice acceptor, BSD = blasticidin, PGK = phosphoglycerate kinase-1 promoter, pA = polyadenylation signal, lacZ-neo = β-galactosidase and neomycin resistance gene. (C) Diagram of the *Mll4* knockout first allele (*Mll4^A^*). The *Mll4* wild type (wt) allele contains 57 coding exons. For abbreviations see Fig. 1B. (D) Loss of MLL4 protein confirmed by Western blot on two *Mll4^A/A^* ESC clones. Immunoblot analysis of extracts from wild type and *Mll4^A/+^* ESC clones detected a signal larger than 460 kDa in agreement with the estimated molecular weight of 600 kDa for MLL4 protein. (E) Dissections from *Mll3^FDC/+^* intercrosses at different stages of development. The homozygous embryos are not distinguishable from their wild type and heterozygous littermates. Scale bar for E9.5 1 mm, for E16.5 2 mm and for P0 2 mm. (F) *Mll4^A/A^* embryos show growth retardation and a visible constriction is formed at the embryonic/extraembryonic boundary (marked by arrowheads). Arrows point at the neural fold. Asterisk marks the allantois. Scale bar 250 μm.

### Mll3 and Mll4 knockouts die at different developmental stages

To explore the function of these conserved proteins, we established multipurpose alleles (Testa et al., 2004) for both *Mll3* and *Mll4* by gene targeting. In our multipurpose allele strategy, frameshifting exons are flanked by loxP sites (exon 49 for *Mll3*; Fig. 1B, Fig. S1A, S1B and exons 2-4 for *Mll4*; Fig. 1C, Fig. S2A, Fig. S2B) accompanied by the insertion of a genetrap stop cassette in the intron upstream of these exons. The stop cassette, which is flanked by FRT sites, contains a *LacZ* reporter and stops target gene transcription because it includes a 5’ splice site, which captures the target gene transcript and a polyadenylation site, which terminates it, thereby - ideally - producing a null allele, termed the ‘*A*’ allele (Testa et al., 2004; Skarnes et al., 2011). After FLP recombination to remove the stop cassette, which establishes the ‘*F*’ allele and restores wild type expression, subsequent Cre recombination establishes a frame-shifted mRNA in the *‘FC’* allele that should provoke nonsense mediated mRNA decay (NMD) (Dyle et al., 2019).

The multipurpose allele strategy aims to establish a loxP allele for conditional mutagenesis and also to mutate the target gene in two different ways, either by truncation of the mRNA (*A* allele) or by NMD (*FC* allele). So if *A/A* and *FC/FC* present the same phenotype, the conclusion that both are null can be established because the *A* and *FC* alleles mutate the gene in different ways. Although unlikely, this conclusion is not secure if the two different mutations produce the same hypomorphic or dominant negative phenotypes.

For *Mll3*, in addition to the multipurpose allele strategy we included a rox-flanked blasticidin-selectable cassette to provide for selection of the 3’ loxP site. Deletion of the rox-flanked cassette by Dre recombinase established the ‘*D’* allele, which is equivalent to the *‘A’* allele described above. Subsequent FLP and Cre recombination establish *‘FD’* and *‘FDC’* alleles that are equivalent to *‘F’* and *‘FC’* alleles respectively.

The multipurpose allele logic worked for *Mll3*. The *D/D* and *FDC/FDC* phenotypes were the same (as described below), supporting the conclusion that the knockout phenotype reported here is the null. In further support of this conclusion, targeting to insert the genetrap cassette into *Mll3* intron 33 presented the same phenotype (Table 1) as those described below for exon 49 targeting. Furthermore, a BayGenomics genetrap in *Mll3* intron 9 also provoked neonatal death (Lee et al., 2013). Although no further analysis other than neonatal death was reported, this outcome concords with the other three *Mll3* mutagenic alleles, supporting the conclusion that MLL3 is not required until birth.

**Table 1.** Genotyping of progeny from Mll3 intercrosses

| **MII3<sup>D/+</sup> x MII3<sup>D/+</sup> (exon 49 floxed)** |  |  |  |  |  |
| --- | --- | --- | --- | --- | --- |
| **Age** | **+/+** | **D/+** | **D/D** | **Total**<br>**(n = litter)** | **Litter size** |
| **P0** | 18.75% | 56.25% | 25%<sup>†</sup> | 16 (2) | 8.0 |
| **E18.5** | 25.3% | 43.4% | 31.3% | 77 (10) | 7.5 |
| **E17.5** | 10.0% | 80.0% | 10.0% | 10 (1) | 10.0 |
| **E16.5** | 10.0% | 70.0% | 20.0% | 10 (1) | 10.0 |
| **E13.5** | 25.0% | 62.5% | 12.5% | 8 (1) | 8.0 |
| **E11.5** | 42.9% | 57.1% | 0.0% | 7 (1) | 7.0 |
| **E10.5** | 17.7% | 64.7% | 17.7% | 17 (2) | 8.5 |
| **E7.5** | 11.1% | 44.4% | 44.4% | 18 (2) | 9.0 |
| **MII3<sup>FDC/+</sup> x MII3<sup>FDC/+</sup> (exon 49 floxed)** |  |  |  |  |  |
| **Age** | **+/+** | **FDC/+** | **FDC/FDC** | **Total**<br>**(n = litter)** | **Litter size** |
| **P0** | 33.3% | 55.6% | 11.1%<sup>†</sup> | 18 (2) | 9.0 |
| **E18.5** | 20.0% | 56.9% | 23.1% | 65 (9) | 7.2 |
| **E17.5** | 17.6% | 58.8% | 23.5% | 17 (2) | 8.5 |
| **E16.5** | 11.1% | 66.7% | 22.2% | 9 (1) | 9.0 |
| **E13.5** | 22.2% | 44.4% | 33.3% | 18 (2) | 9.0 |
| **E11.5** | 11.1% | 66.7% | 22.2% | 9 (1) | 9.0 |
| **E10.5** | 50.0% | 50.0% | 0.0% | 8 (1) | 8.0 |
| **E9.5** | 22.2% | 44.4% | 33.3% | 9 (1) | 9.0 |
| **MII3<sup>D/+</sup> x MII3<sup>D/+</sup> (exon 34 floxed)** |  |  |  |  |  |
| **Age** | **+/+** | **D/+** | **D/D** | **Total**<br>**(n = litter)** | **Litter size** |
| **P0** | 37.5% | 56.3% | 6.3%<sup>†</sup> | 16 (2) | 8.0 |
| **E18.5** | 17.6% | 76.5% | 5.9% | 17 (3) | 5.7 |
| **E17.5** | 25% | 62.5% | 12.5% | 10 (1) | 10.0 |
| **E15.5** | 28.6% | 42.9% | 28.6% | 7 (1) | 7.0 |
| **E13.5** | 25.0% | 50% | 25.0% | 8 (1) | 8.0 |

<sup>†</sup> indicates dead fetuses

For *Mll4*, the *Mll4^A/A^* and *Mll4^FC/FC^* phenotypes were different, potentially revealing different aspects of MLL4 function. As described below, *Mll4^A/A^* embryos are defective before gastrulation whereas *Mll4^FC/FC^* died at birth. As expected, MLL4 expression was abolished by the intron 1 genetrap cassette insertion in *Mll4^A/A^* ESCs (Fig. 1D), indicating that the stronger phenotype is the null. However, mRNA and truncated protein expression persisted in *Mll4^FC/FC^* ESCs (data not shown) indicating that the milder phenotype was hypomorphic. Analysis of this hypomorphic allele will not be presented in this paper. Our conclusions about the null *Mll4* phenotype are supported by a briefly described homozygous BayGenomics *Mll4* genetrap in intron 19 (Lee et al., 2013), which also resulted in embryonic lethality before E10.5. Although not investigated, this phenotype was severe and could be the same as *Mll4^A/A^*. Notably another *Mll4* allele may also present the same phenotype. Aiming to mutate the methyltransferase activity of MLL4, Jang et al. (2017) mutated three conserved tyrosines to alanines in the SET domain. However they observed MLL4 protein instability and may have inadvertently created a null. Again the embryos were not investigated except for observing severe early embryonic lethality that could be the same as *Mll4^A/A^*.

The *Mll3* and *Mll4* null phenotypes are strikingly different. Embryos lacking MLL3 appeared to develop normally until birth whereupon they died because they failed to breathe, although they gasped (Fig. 1E, Table 1). Embryos lacking MLL4 showed abnormalities during gastrulation and died shortly afterwards (Fig. 1F, Table 2). The earliest observable phenotype in *Mll4^A/A^* embryos was growth retardation starting at E6.5 followed by the appearance of a marked constriction at the embryonic/extraembryonic boundary one day later (Fig. 1F). Later in development the *Mll4^A/A^* embryos displayed abnormal headfolds, absence of somites and heart beat, and did not turn (Fig. 1F, Table 2). Despite this severe phenotype, *Mll4^A/A^* embryos displayed no observable change in global mono-, di- or tri-methylation of H3K4 (Fig. S3). Furthermore, heterozygous *Mll4^A/+^* embryos exhibited abnormalities whereas *Mll3^A/+^* embryos were normal.

**Table 2.** Embryonic lethality of Mll4^A/A^ embryos.

| MII4<sup>A/+</sup> X MII4<sup>A/+</sup> |  |  |  |  |  |  |  |
| --- | --- | --- | --- | --- | --- | --- | --- |
| Age | +/+ | A/+ | A/A | Resorbed | Exen-<br>cephaly | Total<br>(n = litter) | Litter<br>size |
| Weaned | 59.1 % | 40.9 % | 0.0 % |  |  | 22 (8) | 2.8 |
| E11.5 | 10.8 % | 48.6 % | 0.0 % | 40.5 % | 10.8 % | 37 (4) | 9.3 |
| E10.5 | 21.5 % | 44.6 % | 13.8 %<sup>†</sup> | 20.0% | 7.7 % | 65 (6) | 10.8 |
| E9.5 | 23.0 % | 45.9 % | 16.2 %<sup>†</sup> | 14.9 % | 9.5 % | 148 (14) | 10.6 |
| E8.5 | 22.9 % | 45.9 % | 17.8 % | 13.4 % | - | 314 (27) | 11.6 |
| E7.5 | 22.1 % | 46.3 % | 18.9 % | 12.6 % | - | 95 (7) | 13.6 |
| E6.5 | 18.8 % | 50.0 % | 12.5 % | 18.8 % | - | 16 (2) | 8.0 |

<sup>†</sup> indicates dead embryos

### Loss of MLL3 results in respiratory failure at birth

Both *Mll3^D/D^* and *Mll3^FDC/FDC^* mice died at birth and were indistinguishable (Table 1). Hence we now term both *Mll3^KO^*. To determine the cause of lethality of *Mll3^KO^* neonates, we followed natural delivery paying attention not to disturb maternal care. *Mll3^KO^* neonates quickly became cyanotic and died immediately after birth or were found dead (Fig. 1E, Table 1). These neonates had normal weight and morphology. The hearts of E15.5 and E18.5 *Mll3^KO^* fetuses showed no morphological abnormalities and were beating indicating normal fetal circulation (Fig. S4A). Neonatal death by asphyxiation can be caused by a defect of the respiratory rhythm generator in the brain stem, which comprises the retrotrapezoïd nucleus/parafacial respiratory group (RTN/pFRG) and pattern generator neurons of the ventrolateral medulla named the pre-Bötzinger complex (preBötC). Knockout of *Jmjd3*, which together with the UTX and UTY subunits of the MLL3/4 complexes are the three known H3K27 demethylases (Van der Meulen et al., 2014), die at birth because the preBötC is absent (Burgold et al., 2012). Therefore we looked for, and found, the preBötC in *Mll3^KO^* perinatal brain stem (Fig. S4B), excluding this explanation for the failure to breathe. Furthermore, the rib cage and palate were intact and the intercostal muscles as well as the diaphragm were normal in *Mll3^KO^* and littermate fetuses (Fig. S4C and Fig. S4D; data not shown).

Considering that the structures of palate, diaphragm, intercostal muscles, brain stem and heart were normal, we focused on defects in the lung as the cause of respiratory failure. In mouse lung development the pseudoglandular stage (E10.5-E16.5) is characterized by dichotomous branching of the bronchi resulting in smaller bronchioles in the periphery. The distal tips form terminal saccules at the cannalicular stage (E16.5-17.5). In the saccular stage (E17.5-P5), the alveolar epithelium differentiates into type I and type II alveolar epithelial cells (Bird et al., 2015).

Macroscopic analysis of *Mll3^KO^* E18.5 lungs revealed mostly a normal morphology with five lobes. Several *Mll3^KO^* E18.5 fetuses, but not all, had smaller lungs than wild type littermates (Fig. 2A). Hematoxylin and eosin staining showed normal proximal conducting airways (Fig. 2A). Staining with the epithelial marker E-cadherin showed normally developed epithelial lining and the integrity of the epithelium was confirmed by the tight junctional marker ZO-1 (Fig. 2B). The differentiated proximal epithelium consists of neuroendocrine, ciliated and secretory (Clara) cells. Cc10, which is a marker for Clara cells, was reduced in both immunofluorescence staining and qRT-PCR in *Mll3^KO^* lungs (Fig. 2B, Fig. 2D). At E18.5 *Mll3^KO^* lungs had thickened alveolar walls and smaller alveolar spaces (Fig. 2A) whereas E15.5 lung morphology was normal (Fig. S5A), suggesting that MLL3 is required for alveolar formation in the saccular phase of lung development. In this phase columnar distal type II alveolar epithelial cells differentiate and give rise to squamous type I alveolar epithelial cells, which line the alveoli and mediate gas exchange (Treutlein et al., 2016). Type II alveolar epithelial cells produce surfactant consisting of phospholipids and the surfactant proteins (SPs) SP-A, SP-B, SP-C and SP-D, which reduce surface tension at the air-liquid interface and prevent alveolar collapse (Woik et al., 2014). In *Mll3^KO^* E18.5 lungs transcripts for the surfactant proteins were lower than in control littermates (Fig. 2D). mRNA levels for Abca3 (ATP-binding cassette sub-family A (ABC1), member 3) which encodes a factor required for lamellar body synthesis in type II alveolar epithelial cells showed a slight increase in *Mll3^KO^* lungs (Fig. 2D). Lysozyme, another marker for type II alveolar epithelial cells, remained unchanged in immunofluorescence staining (Fig. 2C). Aquaporin 5, a characteristic marker for type I alveolar epithelial cells was reduced at both transcript and protein levels (Fig. 2C, Fig. 2D). From these data we conclude a defect(s) in alveolar epithelial cell differentiation and maturation contributed to the respiratory failure.

**Fig. 2.**
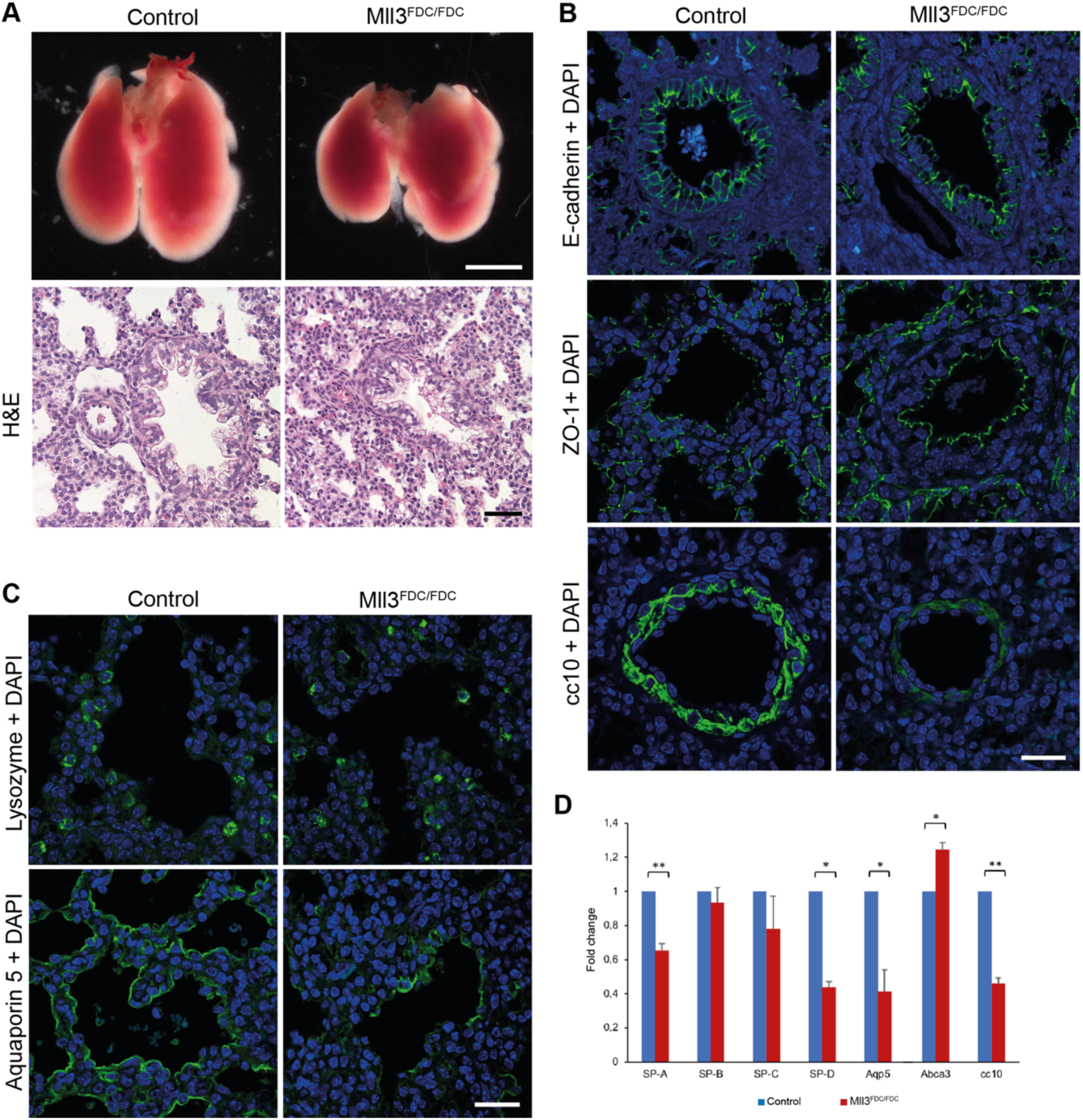
Loss of MLL3 causes defects in lung development. (A) Analysis of gross morphology and H&E staining of *Mll3^KO^* and control littermate lung at E18.5. Scale bar 500 μm (whole lung) and 250 μm (histology). (B) E-cadherin expression in the proximal lung epithelium and ZO-1, a tight junction marker, remain unchanged in control and *Mll3^KO^* littermates. However, expression of cc10 is reduced in *Mll3^KO^* when compared to control littermates. Scale bar 25 μm. (C) Lysozyme expression, a characteristic marker for type II alveolar epithelial cells is comparable in *Mll3^KO^* and control littermate lung. Expression of Aquaporin 5, a marker for type I alveolar epithelial cells is reduced in *Mll3^KO^* when compared to control littermates. Scale bar 25 μm. (D) qRT-PCR analysis of proximal and distal airway cell type markers between control and *Mll3^KO^* lungs at E18.5. Expression levels were normalized to *Rpl19* and plotted as fold change relative to control. Mean +s.d. is shown (*P <0.05 and **P<0.01 as calculated by unpaired t-test).

The specification of the various lung epithelial cell types requires a balance between differentiation and proliferation (Bellusci et al., 1997). The developing lung at the pseudogandular stage is characterized by high proliferative activity, which drastically decreases during sacculation (Fig. S5B). Lungs of both genotypes underwent this inhibition of proliferation however to a lesser extent in the *Mll3^KO^* that correlated with increased septum thickness (Fig. S5B). Concomitantly extracellular matrix (ECM) proteins laminin α1 (Lama1) and fibronectin were apparently more prevalent in the ECM of the basement membrane in *Mll3^KO^* lung (Fig. S5C).

Because MLL3 expression was detected in close proximity to pulmonary vessels (Fig. S6A) and several *Mll3^KO^* lungs were paler than wildtype controls, we examined whether loss of MLL3 impaired pulmonary circulation via improper development of the lung vasculature. However, staining with von Willebrand Factor (vWF) on *Mll3^KO^* lung sections revealed intact endothelium and staining the smooth muscle surrounding the blood vessels with alpha smooth muscle actin (αSMA) was also apparently normal (Fig. S6B). We conclude vascular development was unaffected in lungs of *Mll3^KO^* mice.

### Incompletely penetrant neural tube Mll4 haploinsufficiency with sex distortion

The null alleles of *Mll3* and *Mll4* presented very different phenotypes in another way. No impact of the heterozygous *Mll3* knockout was observed (Table 1), whereas the heterozygous *Mll4* knockout presented a phenotype. Neural fold defects were observed in *Mll4^A/+^* embryos from E9.5 and further throughout gestation. At E9.5, some *Mll4^A/+^* embryos had failed to close their neural tube. Later in embryogenesis from E10.5 until E16.5, exencephaly was observed with open cranial neural folds displaying an everted and enlarged appearance (Fig. 3A, Fig. 3B). This disorder was inherited with a parent-of-origin distortion. When only the father was carrier of the *Mll4^A^* allele, 11% of the heterozygous embryos developed exencephaly, while 50% of the heterozygous embryos were affected if only the mother passed on the allele (Table 3). As expected, when both parents were heterozygous the number of exencephalic heterozygous embryos was in between these two frequencies at 20% (Table 3). Moreover, two-thirds of all exencephalic embryos were female. Later in gestation, exposed neural folds degenerated, producing anencephaly (data not shown). Embryos with such disorder were found dead at P1 as they were either stillborn or died within the first hours after birth, hence the low recovery rate of *Mll4^A/+^* pups (Table 3).

**Table 3.** Exencephaly of Mll4^A/+^ embryos is inherited in a parent-of-origin dependent manner

| ♂ MII4<sup>A/+</sup> X ♀ CD1 |  |  |  |  |  |  |  |
| --- | --- | --- | --- | --- | --- | --- | --- |
| Age | +/+ | A/+ | A/A | Resorbed | Exen-<br>cephaly | Total<br>(n = litter) | Litter<br>size |
| Weaned | 82.1 % | 17.9 % |  |  |  | 1508 (181) | 8.3 |
| E18.5 | 53.7 % | 46.3 % | 0.0 % | 0.0 % | 12.0 % | 54 (4) | 13.5 |
| E16.5 | 40.0 % | 60.0 % | 0.0 % | 0.0 % | 22.2 % | 15 (1) | 15.0 |
| E15.5 | 46.2 % | 50.0 % | 0.0 % | 3.8 % | 0.0 % | 26 (2) | 13.0 |
| E14.5 | 75.0 % | 25.0 % | 0.0 % | 0.0 % | 0.0 % | 12 (1) | 12.0 |
| E13.5 | 43.9 % | 54.4 % | 0.0 % | 1.8 % | 12.9 % | 57 (4) | 14.3 |
| E11.5 | 47.1 % | 52.9 % | 0.0 % | 0.0 % | 11.1 % | 17 (1) | 17.0 |
| E10.5 | 51.2 % | 46.4 % | 0.0 % | 2.4 % | 17.9 % | 84 (6) | 14.0 |
| E9.5 | 44.8 % | 51.2 % | 0.0 % | 4.0 % | 7.8 % | 125 (9) | 13.9 |
| E8.5 | 23.1 % | 76.9 % | 0.0 % | 0.0 % | - | 13 (1) | 13.0 |

| ♂ CD1 X ♀ MII4<sup>A/+</sup> |  |  |  |  |  |  |  |
| --- | --- | --- | --- | --- | --- | --- | --- |
| Age | +/+ | A/+ | A/A | Resorbed | Exen-<br>cephaly | Total<br>(n = litter) | Litter<br>size |
| Weaned | 77.2 % | 22.8 % | 0.0 % |  |  | 114 (20) | 5.7 |
| E11.5 | 58.3 % | 25.0 % | 0.0 % | 16.7 % | 66.7 % | 12 (1) | 12.0 |
| E10.5 | 28.0 % | 52.0 % | 0.0 % | 20.0 % | 30.8 % | 25 (2) | 12.5 |
| E9.5 | 36.8 % | 42.1 % | 0.0 % | 21.1 % | 75.0 % | 19 (2) | 9.5 |

**Fig. 3.**
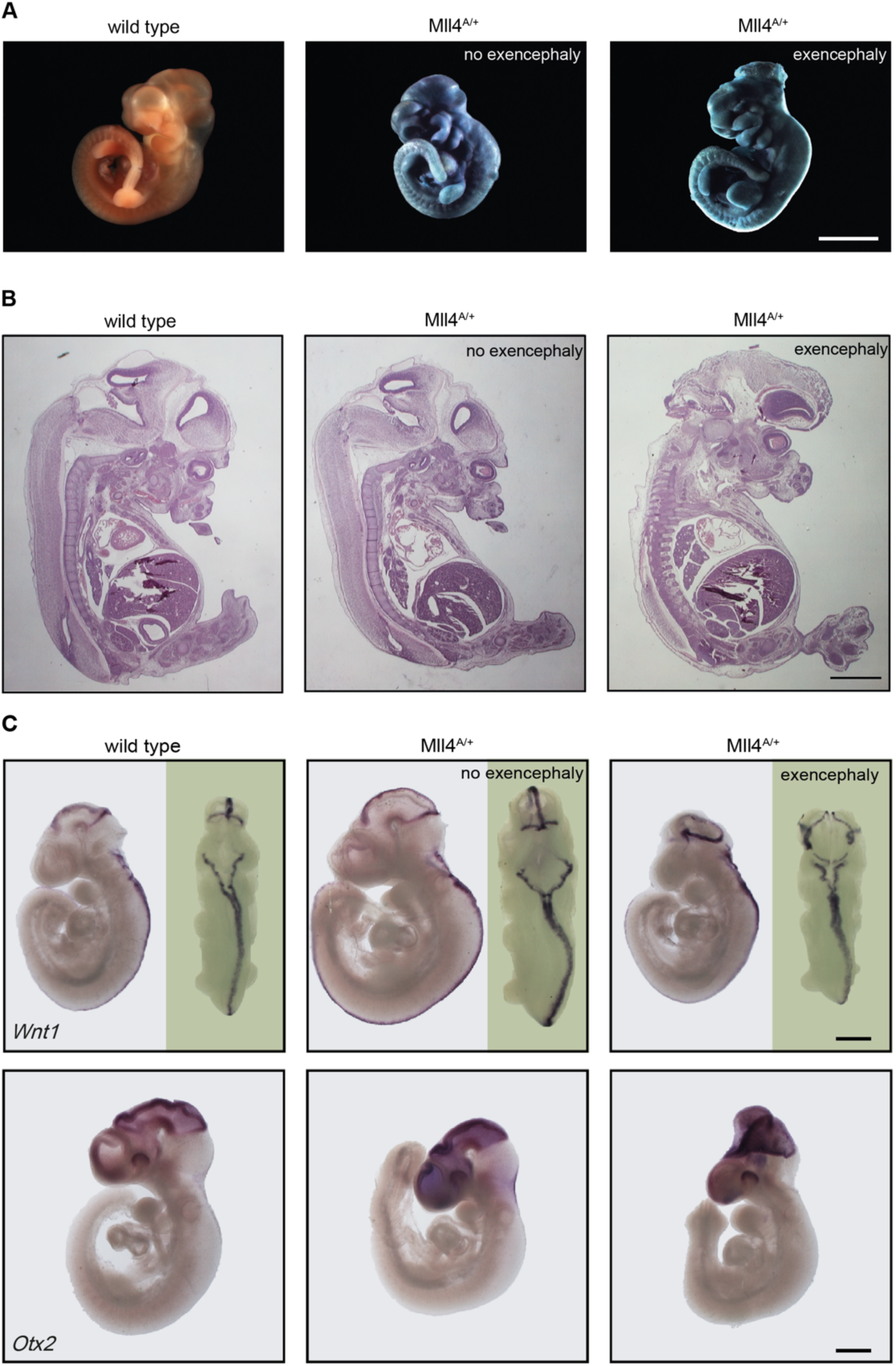
Neural tube closure defects in *Mll4^A/+^* embryos. (A) β-galactosidase staining of heterozygous *Mll4^A/+^* embryos at E10.5. The embryo on the right exhibits exencephaly. Scale bar 1 mm. (B) H&E-stained sagittal sections of E13.5 embryos show the everted and enlarged exposed neural folds. Scale bar 1 mm. (C) *In situ* hybridization analysis for *Otx2* and *Wnt1* expression at E9.5. The expression of these markers was not affected in exencephalic *Mll4^A/+^* embryos compared to wild type littermates. Scale bar 250 μm.

Because the *Wnt1* gene is only 40 kb 3’ of the *Mll4* gene on mouse chromosome 15, and *Wnt1* is typically expressed in the dorsal midline of the developing hindbrain and spinal cord (Parr et al., 1993), we analyzed its’ expression using *in situ* hybridization. As seen in Figure 3C, *Wnt1* is expressed normally in the exencephalic embryo so disturbed *Wnt1* expression is not the cause of the phenotype. We also evaluated *Otx2* expression, which is located to the forebrain and midbrain and characterized by a sharp mid-hindbrain boundary (Ang et al., 1994). It was also unaffected by exencephaly (Fig. 3C).

### Adult heterozygous Mll4 mice present aspects of Kabuki syndrome

In humans, all *Mll4* mutations associated with Kabuki syndrome are heterozygous. Having observed an embryonic heterozygous phenotype, we therefore looked for signs of Kabuki syndrome in viable *Mll4^A/+^* mice after birth. Surviving *Mll4^A/+^* pups were analyzed for facial, cranial and skeletal abnormalities but none were observed. However we did find indications of the metabolic problems that are associated with Kabuki syndrome. Significant differences in body weight between wild type and *Mll4^A/+^* mice of both sexes from the age of four weeks to 21 weeks could be observed. *Mll4^A/+^* mice of both sexes remained 30% smaller throughout adulthood (Fig. 4A) and showed a decreased amount of white adipose tissue (data not shown). Glucose and insulin tolerance tests were performed in reaction to the decreased body weight of the heterozygous mice *Mll4^A/+^*. Although *Mll4^A/+^* mice had the same fasting blood glucose levels as wild type littermates, male *Mll4^A/+^* mice displayed altered glucose tolerance as their blood glucose levels declined faster and reached their initial values earlier (Fig. 4B). Also when performing insulin tolerance tests male *Mll4^A/+^* mice showed higher insulin sensitivity compared to wild type mice (Fig. 4C). In contrast female *Mll4^A/+^* mice displayed a slight impairment in glucose tolerance and no change in insulin tolerance tests (Fig. 4B, Fig. 4C).

**Fig. 4.**
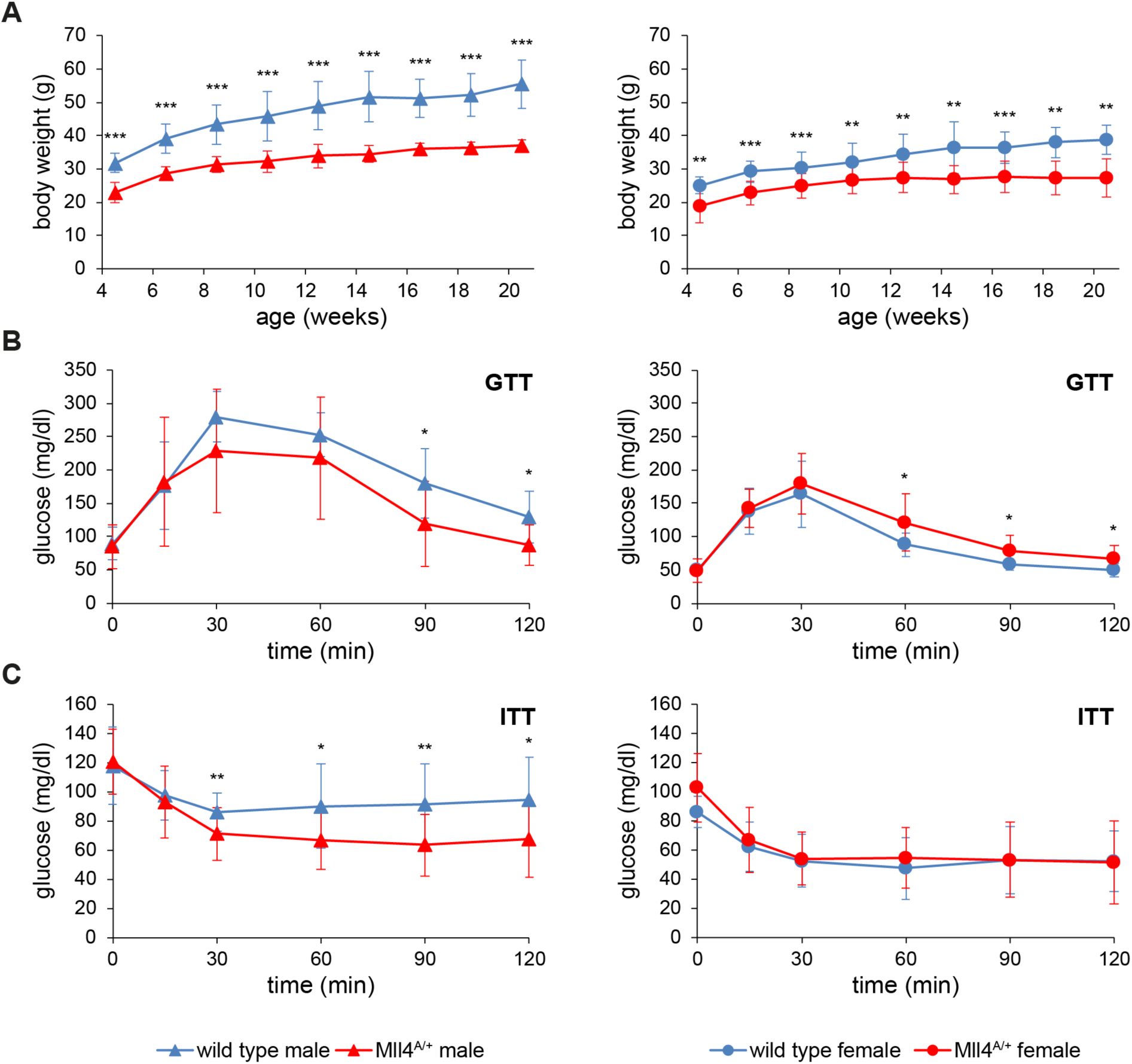
Decreased body weight and hypoglycemia of *Mll4^A/+^* mice. (A) Development of body weight over 4-20 weeks for a total of 48 mice (24 males: 15 wild type and 9 heterozygous mice; 24 females: 16 wild type and 8 heterozygous mice) is shown. The differences in weight are significant at any developmental stage. Mean ± s.d. is shown (**P<0.01 and ***p <0.001 as calculated by unpaired t-test). (B) For the glucose tolerance test (GTT), in total 30 mice (14 males: 7 wild type and 7 heterozygous mice; 16 females: 8 wild type and 8 heterozygous mice) at an age of 9 to 15 weeks have fasted for 16 h, until a glucose solution (1.5 mg/g body weight) was orally applied. Blood glucose levels were measured over a period of 120 min. Mean ± s.d. is shown (*P<0.05 as calculated by unpaired t-test). (C) Insulin tolerance tests (ITT) after 6 h of fasting were performed with 45 mice (30 males: 14 wild type and 16 heterozygous mice; 15 females: 7 wild type and 8 heterozygous mice) between the age of 12 and 45 weeks. Insulin (0.75 mU/g body weight) was injected intraperitoneally. Blood glucose levels were measured within a time frame of 120 min. Mean ± s.d. is shown (*P<0.05 and **p <0.01 as calculated by unpaired t-test).

### Mll4 is required for specification of the embryonic anterior-posterior axis

In contrast to the incompletely penetrant heterozygous *Mll4* knockout, all *Mll4^A/A^* embryos displayed a phenotype similar to that observed in knockouts of *Hnf3β* (*Foxa2*), *Otx2* and *Lim1* (*Lhx1*), in which specification of the anterior-posterior (A-P) embryonic axis is disrupted (Kinder et al., 2001). To examine A-P patterning in *Mll4^A/A^* embryos, we performed whole-mount *in situ* hybridizations for anterior and posterior molecular markers (Fig. 5, Fig. 6) between developmental stages E6.5 to E7.75. *Lefty1*, an early anterior marker expressed at the anterior visceral endoderm (AVE) of E6.5 embryos was absent in *Mll4^A/A^* embryos. Expression of another AVE marker, *Hex,* was restricted to the distal region of *Mll4^A/A^* embryos. *Otx2* was prominent at the anterior ectoderm and visceral endoderm of control embryos but was detected at the distal epiblast and the overlying visceral endoderm of *Mll4^A/A^* embryos (Fig. 5A). *Cerberus1* (*Cer1*), a Bmp4, Nodal and Wnt antagonist is normally expressed at the AVE (Piccolo et al., 1999) but only weakly expressed at the distal region in *Mll4^A/A^* embryos (Fig. 5B). The Wnt antagonist, *Dkk1*, is typically expressed closest to the embryonic/extraembryonic boundary. However, in *Mll4^A/A^* embryos expression was observed at the distal region (Fig. 5C).

**Fig. 5.**
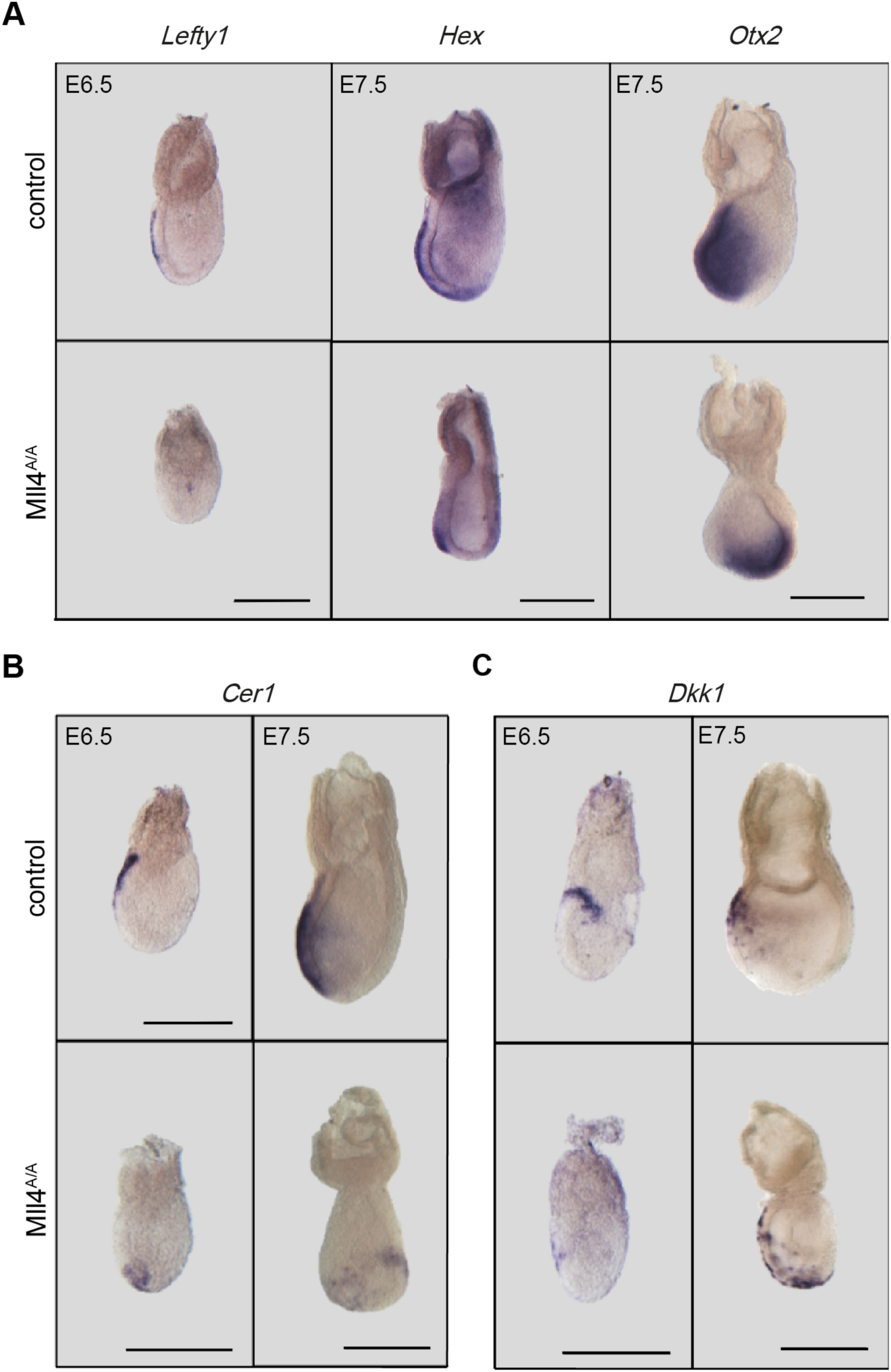
*Mll4^A/A^* embryos show defective patterning of the AVE from E6.5 to E7.5. (A) *Lefty1* (n=5/5), *Hex* (n=5/6), *Otx2* (n=2/3) expression in control and *Mll4^A/A^* embryos between E6.5 and E7.75. All embryos are oriented with the anterior to the left. Scale bar 250 μm. (B) (C) *Cer1* (n=4/4 for E6.5 and n=5/5 for E7.5) and *Dkk1* (n=2/2 for E6.5 and n=2/4 for E7.5) expression in control and *Mll4^A/A^* embryos between E6.5 and E7.5. All embryos are oriented with the anterior to the left. Scale bar 250 μm.

**Fig. 6.**
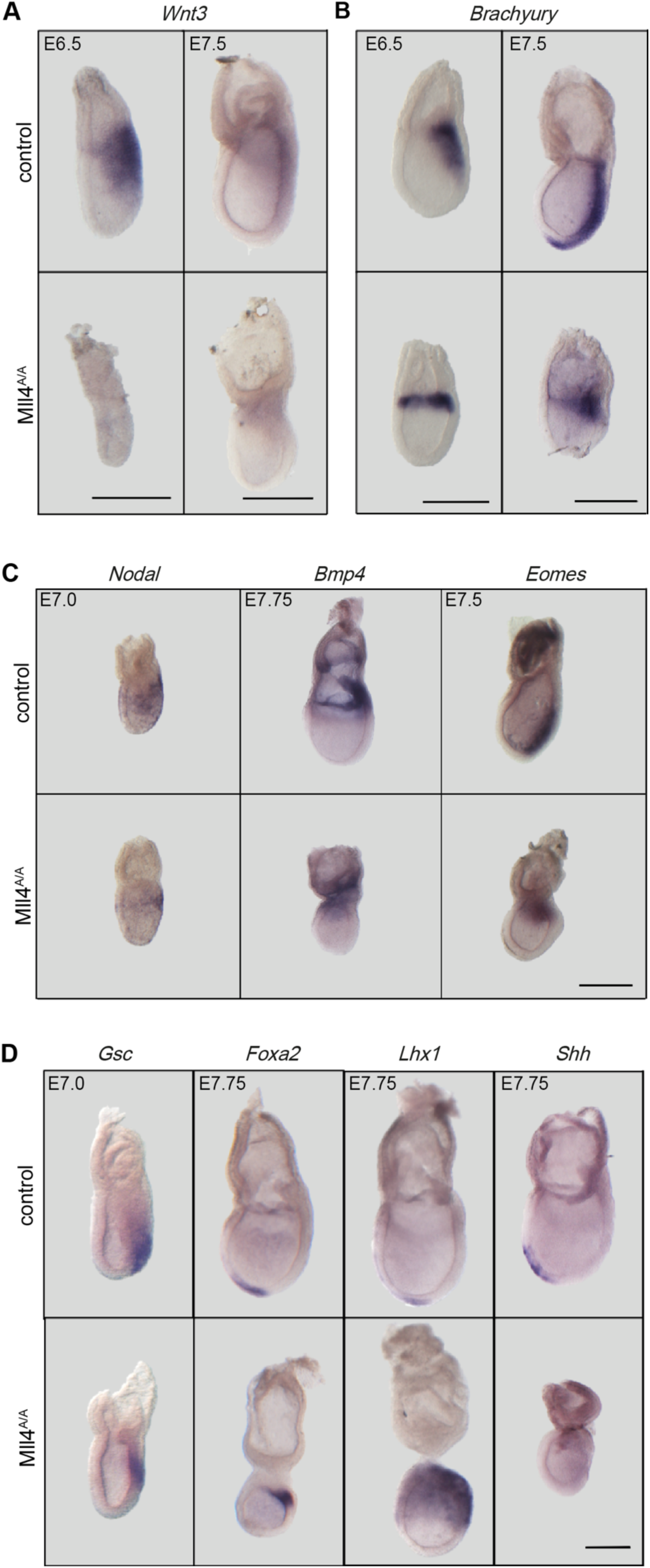
Ectopic expression of primitive streak and node markers in *Mll4^A/A^* embryos. (A) (B) *Wnt3* (n=1/1 for E6.5 and n=2/4 for E7.5) and *Brachyury* (n=1/1 for E6.5 and n=3/3 for E7.5) marks the length of primitive streak (PS) as it forms in control embryos from E6.5 to E7.5, but in *Mll4^A/A^* embryos they are expressed at the embryonic/extraembryonic boundary and do not extend till the distal tip. All embryos are oriented with the anterior to the left. Scale bar 250 μm. (C) In control embryos, *Nodal* (n=4/7)*, Bmp4* (n=3/3) and *Eomes* (n=6/7) are expressed at the PS, posterior PS and anterior PS, respectively. In *Mll4^A/A^* embryos the expression is restricted to the embryonic/extraembryonic boundary. All embryos are oriented with the anterior to the left. Scale bar 250 μm. (D) *Gsc* (n=2/2)*, Foxa2* (n=3/4) and *Lhx1* (n=3/4) are expressed at the node and node derivatives of control embryos but are proximally expressed in *Mll4^A/A^* embryos. The node derivative marker *Shh* (n=3/3) is completely absent in *Mll4^A/A^* embryos. All embryos are oriented with the anterior to the left. Scale bar 250 μm.

Since the anterior markers were misexpressed at the distal region of *Mll4^A/A^* embryos (Fig. 5), we expected defective posterior patterning. *Wnt3* and *Brachyury* are the earliest markers of the primitive streak confined exclusively to the posterior side of control embryos. However, in *Mll4^A/A^* embryos the expression was seen ectopically at the embryonic/extraembryonic boundary (Fig. 6A, Fig. 6B). *Nodal* is essential for the induction and maintenance of the primitive streak (Conlon et al., 1994). Normally *Nodal* is expressed in the posterior embryonic ectoderm, expanding along the length of the primitive streak. *Nodal* did not extend as far distally in *Mll4^A/A^* embryos. In control embryos, *Bmp4* and *Eomes* marked the posterior and anterior region of the primitive streak, respectively. In addition, they are markers of the extraembryonic chorion and amnion. In *Mll4^A/A^* embryos both markers were detected at the embryonic/extraembryonic boundary (Fig. 6C). The patterning of posterior markers in *Mll4^A/A^* embryos indicates that the primitive streak did not extend distally. Consequently the mesodermal wings were absent (Fig. S7).

At the anterior region of the primitive streak is a distinct region called the node from which the anterior mesendoderm derivatives such as head process and notochord develop (Benazeraf and Pourquie, 2013). Since the primitive streak did not extend till the distal tip of the *Mll4^A/A^* embryos, we examined if the node and the node derivatives were present in *Mll4^A/A^* embryos. *Goosecoid* (*Gsc*) was distally induced in control embryos but expression was more confined and proximal in *Mll4^A/A^* embryos. *Foxa2* and *Lhx1* marked the node and its derivatives in control embryos. In *Mll4^A/A^* embryos expression for both was seen at the proximal epiblast. *Shh* was expressed exclusively in the head process arising from the node in control embryos but was completely absent in *Mll4^A/A^* embryos (Fig. 6D). Despite the fact that *Mll4^A/A^* embryos expressed markers of the node they failed to establish the node derivatives.

At E8.5 *Mll4^A/A^* mutants were characterized by the absence of *Mox1* transcripts confirming the aforementioned lack of somites (Fig. S7). Very weak and discontinuous localization of *Brachyury* verified that the node derivatives were not specified (Fig. S7). We noticed weak expression of *Hoxb1* suggesting the presence of precursor cells for rhombomere 4 (Fig. S7).

### Mll4 is essential for AVE migration

Shortly after implantation, the first asymmetry in normal embryos is evident as a proximal-distal (P-D) axis (Beddington and Robertson, 1999). At the distal tip, AVE cells undergo a transition from columnar to squamous, protrude filipodia and migrate unidirectionally as a collective towards the future anterior of the embryo until they reach the embryonic/extraembryonic boundary (Srinivas et al., 2004). At E6.5 squamous AVE cells form a single-layer reaching the embryonic/extraembryonic boundary indicated by expression of HEX (Fig. 7A). In *Mll4* deficient embryos, AVE cells failed to reach the embryonic/extraembryonic boundary. Notably, HEX expressing cells retained a cuboidal shape and displayed strong apical actin (Fig. 7A) indicating that the first defect in *Mll4^A/A^* embryos is a failure to undergo the columnar to squamous transition that precedes migration.

**Fig. 7.**
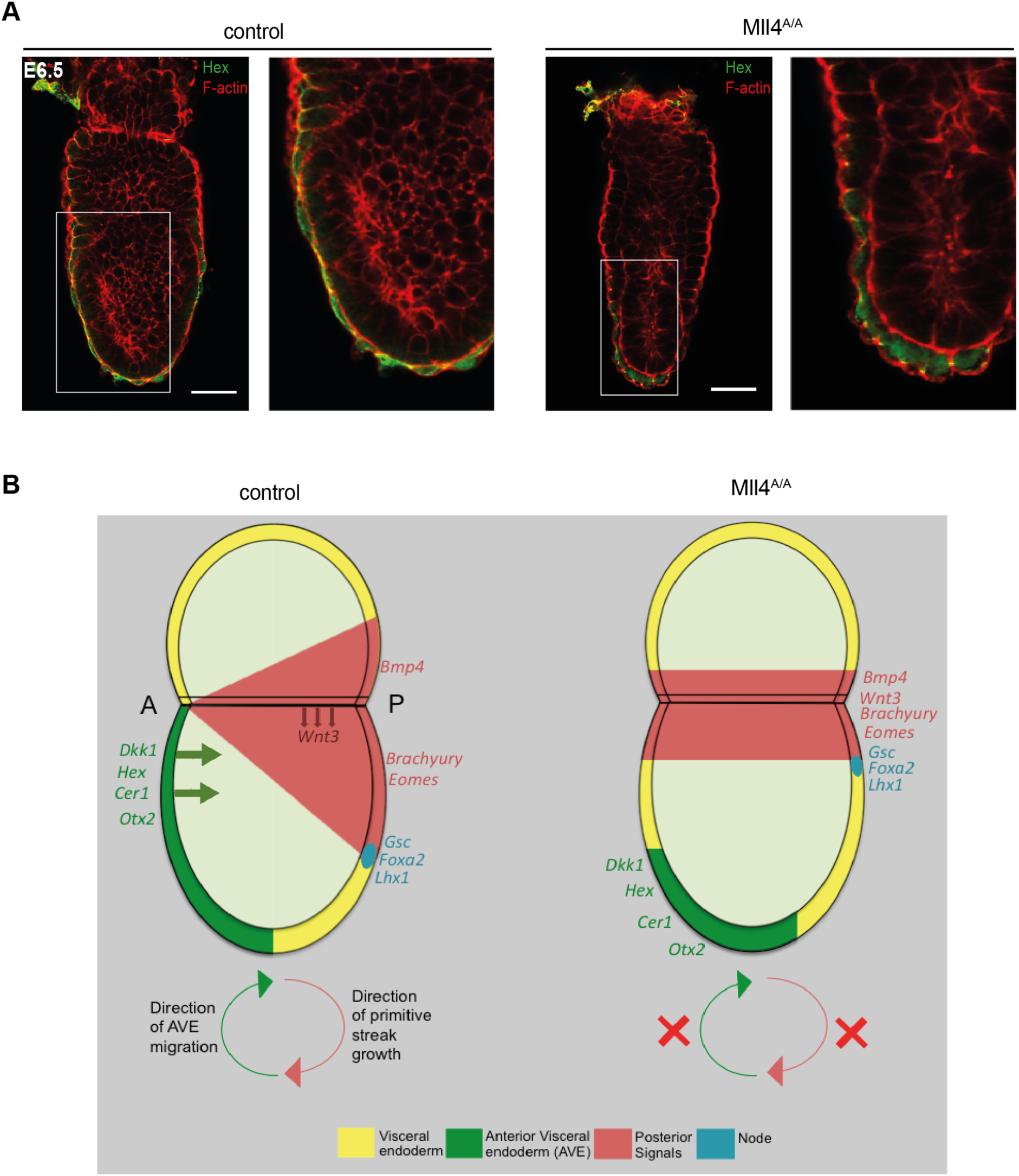
Model for MLL4 function in establishing A-P patterning. (A) Epithelial organization of embryos at E6.5 of control and *Mll4^A/A^* genotype. Individual confocal sections of embryos stained for F-actin (red) and Hex (green) (n=5/5). The boxed region is magnified on the right. Scale bar 50 μm. (B) In control embryos, the AVE (green) migrates to the anterior, restricting the primitive streak to the posterior (red). Thus, the A-P axis is established and gastrulation progresses. In the absence of MLL4, the AVE is at the distal region, causing the primitive streak to form at the embryonic/extraembryonic boundary. The embryo thus undergoes defective gastrulation resulting in embryonic lethality indicating a functional role of MLL4 in AVE migration.

## Discussion

Despite the similarities between MLL3 and MLL4 including protein architecture, residence in apparently identical protein complexes, ubiquitous expression and high frequency of mutations in almost all human cancers, their null phenotypes in mouse development are dramatically different. MLL4 is indispensable for the establishment of the A-P axis and progression of gastrulation. MLL3 is not required until the final steps of lung development to ensure neonatal breath at birth. It therefore appears that these two conserved sister proteins are required for very different, highly specific functions in mouse development. In this regard, they are similar to MLL1, which is first required for definitive hematopoiesis at E12.5 with apparently little other contribution to the developing embryo (Ernst et al., 2004). The observation that these highly conserved proteins are only required for a few, very specific, developmental functions does not concur with the prevalent model that MLL3 and MLL4 are the transcription co-factors that deposit the universal epigenetic characteristic of enhancers, H3K4me1 (Lee et al., 2013; Rao and Dou, 2015; Piunti and Shilatifard, 2016). This conundrum could find a resolution if the MLLs are embedded in comprehensive functional backup and their individual knockout phenotypes only reveal flaws in this comprehensive backup. Some evidence for this proposition has been acquired (Lee et al., 2013; Denissov et al., 2014; Chen et al., 2017; Chen et al., 2018).

### Mll3 and defective respiration

*Mll3^D/D^* and *Mll3^FDC/FDC^* neonates died due to failures in the final steps of lung maturation. MLL3 is not required for patterning of the lung but is required for (i) efficient differentiation of distal lung epithelium when squamous type I alveolar epithelial cells arise from columnar type II alveolar epithelial cells, and (ii) thinning of the mesenchyme possibly due to sustained proliferative activity during the cannalicular/saccular phase. The concomitant occurrence of defects in two nearby cell types suggest an underlining failure of cell signaling during the final maturation of the lung.

### Mll4 and defective AVE migration

Migration of the AVE is the first collective cell migration in the mouse embryo and precedes gastrulation (Rossant and Tam, 2009) (Fig. 7B). Before migration extraembryonic visceral endoderm cells change their shape from columnar to squamous, which involves cytoskeletal rearrangements and the projection of filopodia towards the direction of migration (Srinivas et al., 2004). At the migratory front of the cell the Rho GTPase, RAC1, is positioned to regulate actin polymerization. Notably, the *Rac1* knockout phenotype is comparable to *Mll4^A/A^* characterized by failed AVE migration and lethality before E9.5 (Sugihara et al., 1998; Migeotte et al., 2010). NAP1, a component of the WAVE complex acts downstream of RAC1 to control actin branching and *Nap1* mutants do not establish the A-P axis due to failed AVE polarization and migration (Rakeman and Anderson, 2006). Considering these similarities, we suggest that MLL4 regulates RAC1 and/or the WAVE complex. Another possibility is that MLL4 acts through interaction with UTX, which was found in an unbiased screen to regulate cell migration (Thieme et al., 2013).

In the absence of MLL4, gene expression associated with the AVE including *Hex, Dkk1 and Cer1* was established but mislocalized at the distal region of the embryo due to the absence of migration. This migration is essential for the correct patterning of both the anterior and posterior of the embryo (Stower and Srinivas, 2014). Consequently the failure of the AVE to reach the anterior embryonic/extraembryonic boundary resulted in failed elongation of the primitive streak towards the distal tip indicated by misexpression of *Brachyury*, *Wnt3*, *Bmp4* and *Eomes*. As a result, node markers *Goosecoid*, *Foxa2* and *Lhx1* were restricted to the posterior-proximal region (Fig. 7B). These failures preceded the absence of mesodermal derivatives emerging from the primitive streak

### MLL4 haploinsufficiency

MLL3 and MLL4 also differ regarding their heterozygous phenotypes. In our survey of the six H3K4 methyltransferases in mouse development (Glaser et al., 2006; Glaser et al., 2009; Andreu-Vieyra et al., 2010; Bledau et al., 2014; Denissov et al., 2014; Brici et al., 2017; Chen et al., 2017; Hanna et al., 2018), only the knockout of *Mll4* has presented embryonic haploinsufficiency. Notably *Drosophila Trr* also displays haploinsufficiency (Chauhan et al., 2012). Amongst the various explanations for haploinsufficiency, transcriptional synergy may be relevant for MLL4. Transcriptional synergy is based on co-operative recruitment of transcription factors to cis regulatory elements to achieve a transcriptional output, which involves a threshold and sigmoidal response to protein concentration (Veitia et al., 2018). Furthermore, a contribution by MLL4 to the stability of decisions based on stochastic choices could explain the incomplete penetrance (Cook et al., 1998). Incomplete penetrance in a rate limiting developmental choice pathway implies that MLL4 does not make the choice rather reduces the error rate by either stabilizing the choice or counter-acting mistakes. A primary role for epigenetic regulation in error reduction concords with our recent findings in yeast where Set1C and the H3K4me3 demethylase Jhd2 act together as a quality control mechanism to ensure symmetrically trimethylated nucleosomes (Choudhury et al., 2019).

The frequency of neural tube defects in *Mll4^A/+^* embryos was affected by the sex of the mutant parent. The defects were more likely when the null allele was transmitted from the mother. Thus it seems likely that MLL4 contributes to oogenesis as well as neurulation. Alternatively, the sex distortion may relate to the sex specific difference between MLL4 complexes, which in females includes only UTX whereas in males both UTX and UTY are involved.

More than 300 genes have been implicated in neural tube closure defects, which are very common in humans estimated at 1 per 1000 fetuses (Juriloff and Harris, 2018), and epigenetic mechanisms involving DNA and histone methylation have emerged as particularly important (Harris and Juriloff, 2010). Notably, folic acid supplementation during pregnancy, which elevates S-Adenosyl Methionine (SAM) levels, diminishes the probability of neural tube closure defects in certain cases (Greene and Copp, 2005). The accompanying proposition that females are more susceptible to neural tube defects due to the increased requirement for SAM in X-chromosome inactivation (Juriloff and Harris, 2000), may be relevant to our observations of sex distorted neural tube closure defects.

Concordant with *Mll4* haploinsufficiency in the mouse, *de novo* heterozygous mutations of *Mll4* are the primary cause of the rare congenital Kabuki syndrome, which involves mental retardation and a distinctive facial appearance. Further features of this phenotypically variable disorder include postnatal dwarfism, heart and kidney dysfunction, skeletal abnormalities, loss of hearing, gastrointestinal disorders and metabolic imbalances including hypoglycemia (Banka et al., 2012; Banka et al., 2015; Bogershausen et al., 2015; Yap et al., 2019). A very mild version of Kabuki syndrome has been observed in a mouse model based on a heterozygous, hypomorphic *Mll4* allele (Benjamin et al., 2017). Similarly, the *Mll4^A/+^* phenotype only presents limited aspects of Kabuki syndrome: reduced body weight, stunted growth and hypoglycemia together with reduced body fat. Nevertheless, these haploinsufficient *Mll4* observations indicate that the amount of expressed MLL4 is critical to its function. *Mll4* haploinsufficiency revealed a requirement for MLL4 function in development after its role in the AVE columnar to squamous transition. *Mll4* conditional mutagenesis also revealed later requirements in heart development, myogenesis and adipogenesis (Lee et al., 2013; Ang et al., 2016). Notably, Ang et al (2016) presented evidence of aortic haploinsufficiency.

Because MLL4 is required for the first collective cell migration in mouse development, and because it is required for the cellular migration processes involved in closure of the neural tube, we suggest that MLL4 is a master regulator of cell migration gene expression programs. Although diverse and as yet only partially documented, the evidence for the various MLL4 functions in mouse development are nevertheless still limited to a small group of specific indications. For MLL3, the indications are even more limited and together, these indications do not offer an explanation for the extraordinary prevalence of *Mll3* and *Mll4* mutations in human cancers. The proposition that the MLL system is deeply embedded in functional redundancy and backup may resolve this conundrum. Testing this proposition requires concerted conditional mutagenesis, which is underway.

## Acknowledgements

We thank Mandy Obst, Doris Müller, Isabell Kolbe, Madeleine Walker and Stefanie Weidlich for excellent technical assistance. We also thank the Biomedical Services (BMS) of the Max Planck Institute of Molecular Cell Biology and Genetics, Dresden for the excellent service and technical assistance. We thank Dr. Siddharth Banka (University of Manchester, UK) for discussions and Dr. Tristan A. Rodriguez (Imperial College, London, UK), Dr. Karin Schuster-Gossler (MH Hannover, Germany), Dr. Janet Rossant (University of Toronto, Canada), Dr. Elizabeth Robertson (University of Oxford, UK), Dr. Dominic Norris (MRC Harwell, UK), Dr. Rachel D. Mullen (MD Anderson, Texas, USA), Dr. Hans Schöler (Max Planck Institut Münster, Germany) and Dr. Andrew P. McMahon (University of Southern California, USA) for providing probes for whole mount *in situ* hybridization and Dr. Go Shioi (RIKEN Kobe, Japan) for providing the anti-Hex antibody and advice. The Advanced Imaging Facility, a core facility of the CMCB Technology Platform at TU Dresden, http://biotp.tu-dresden.de/facilities/advanced-imaging/ assisted this research.

## Author contributions

D.A., Q.Z., A.S.B., C.M., A.F.S., K.A and A.K. designed and performed experiments, D.A., Q.Z, A.K., A.S.B. and C.M. examined the embryonic phenotype, A.S.B., J.F., K.A. generated DNA reagents, A.K., D.A. and A.F.S. wrote the manuscript.

---

**Fig. S1.**
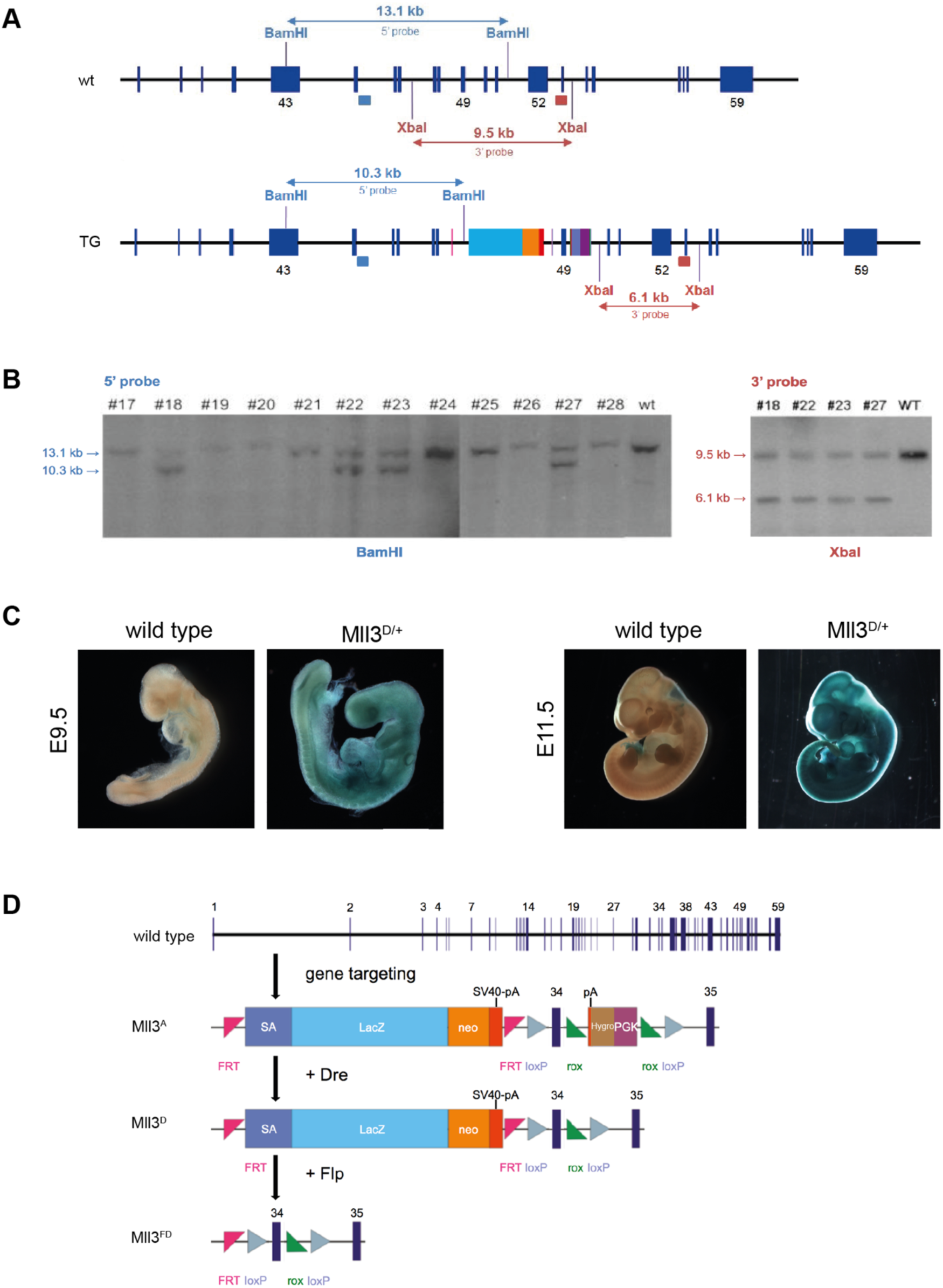
Gene targeting strategy for *Mll3*. (A) Schematic representation of the Southern strategy to identify correctly targeted events in the *Mll3* locus using 5’ (light blue box) and 3’ (red box) external probes. Blue boxes with numbers underneath indicate exons. (B) Southern blot analysis using 5’ and 3’ external probes. Clones #18, #22, #23 and #27 carried the correctly targeted event. (C) Wild type and *Mll3^D/+^* embryos at E9.5 and E11.5 were stained for β-galactosidase. Scale bar 500 μm. (D) Diagram of the *Mll3* knockout first allele (*Mll3^A^*) with floxed exon 34. Instead of the PGK-Blasticidin-polyA a PGK-Hygromycin-PolyA cassette was inserted. For abbreviations see Fig. 1B.

**Fig. S2.**
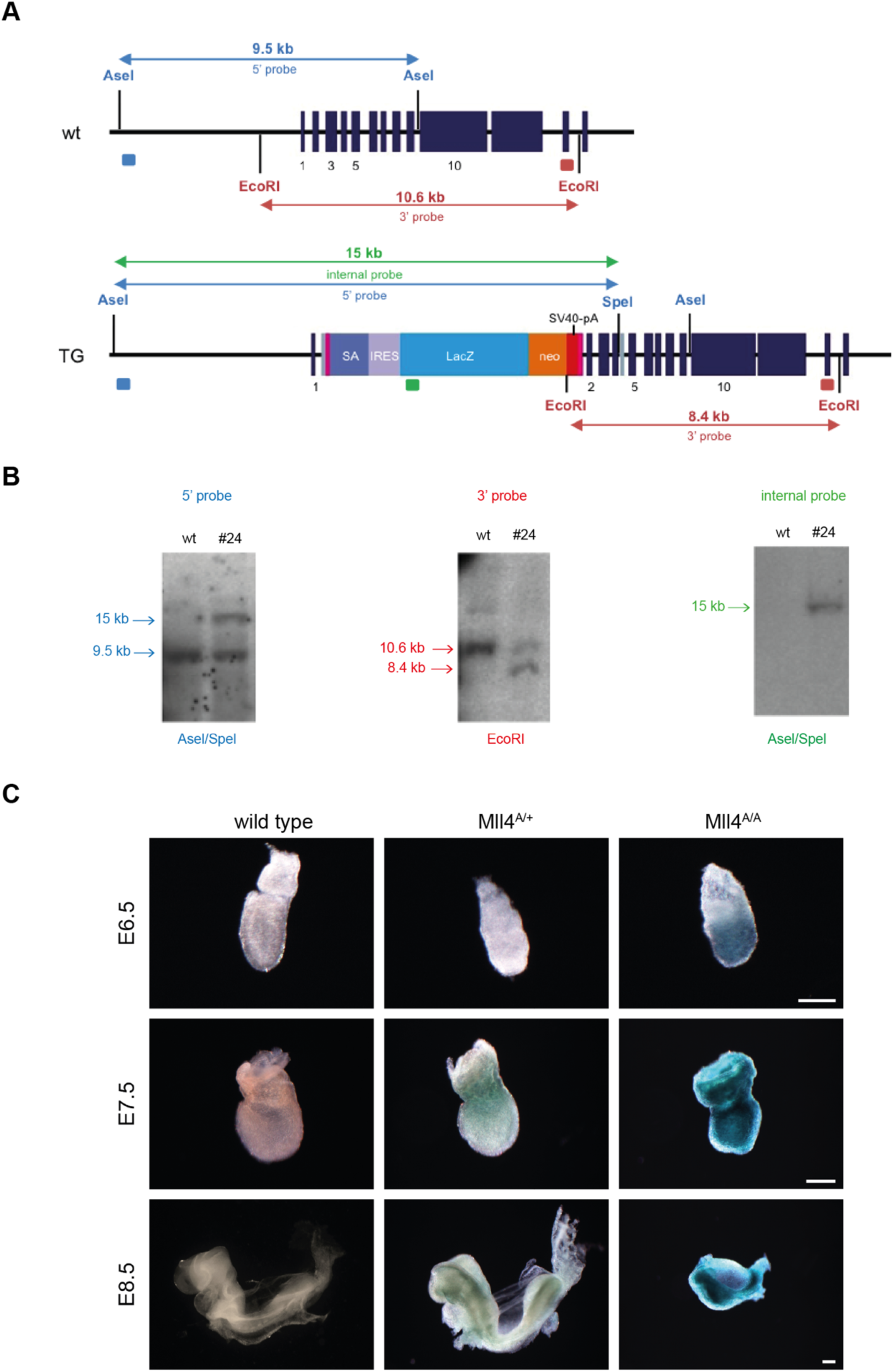
Gene targeting strategy for *Mll4*. (A) Schematic representation of the Southern blot strategy to identify correctly targeted events in the *Mll4* locus using 5’ (light blue box), 3’ (red box) external probes and the internal LacZ (green box) probe. Blue rectangles with numbers underneath indicate exons. (B) Southern blot analysis using 5’ and 3’ external probes and LacZ internal probe. Only clone #24 is shown. (C) Wild type and *Mll4^A/+^* embryos at E6.5, E7.5 and E8.5 were stained for β-galactosidase. Scale bar 250 μM.

**Fig. S3.**
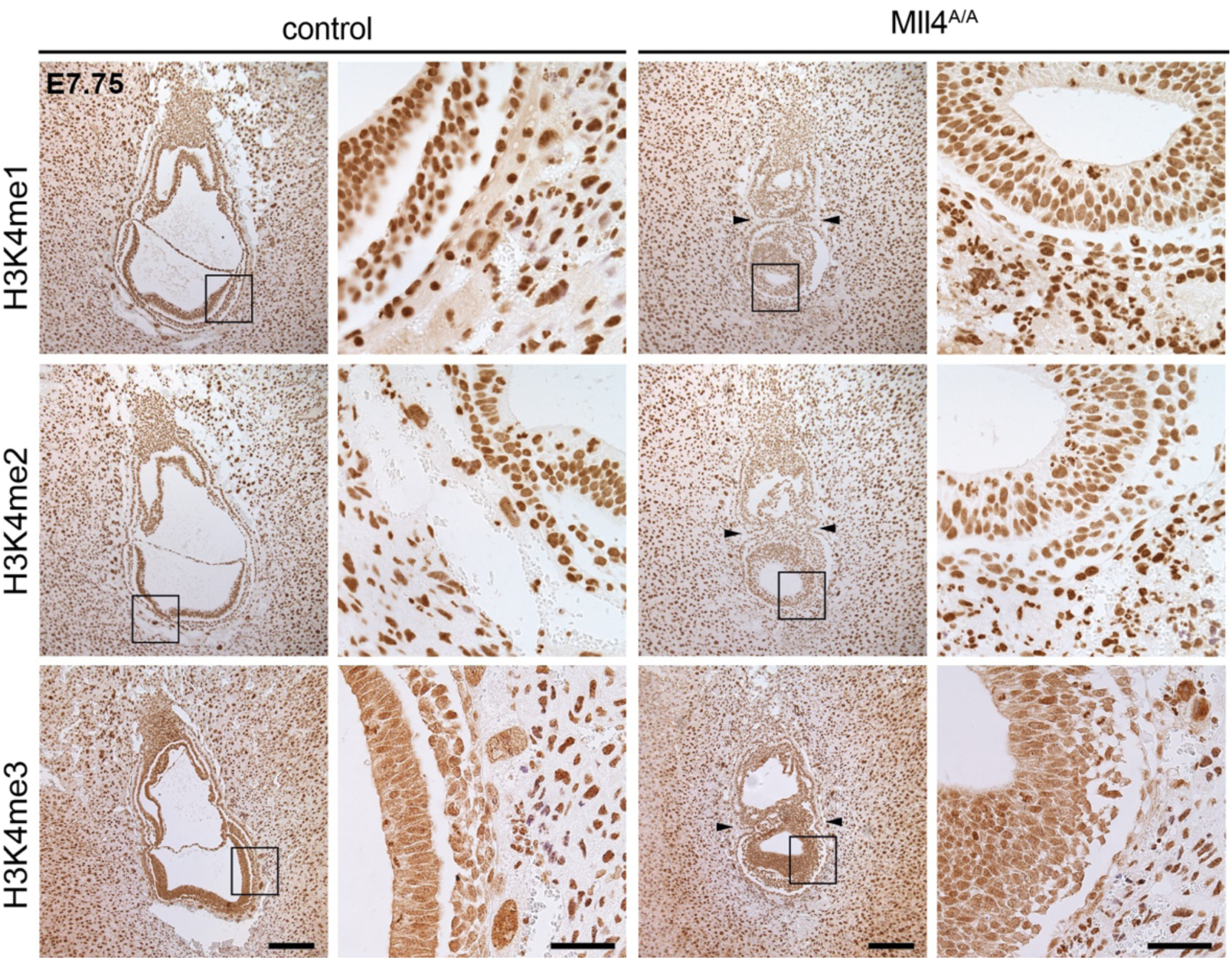
No global changes in H3K4 methylation levels. Immunohistochemistry on sections of wild type and *Mll4^A/A^* E7.75 embryos *in utero* with H3K4me1-, H3K4me2- and H3K4me3-specific antibodies. The boxed region is magnified on the right. Scale bars 50 μm and 200 μm, respectively.

**Fig. S4.**
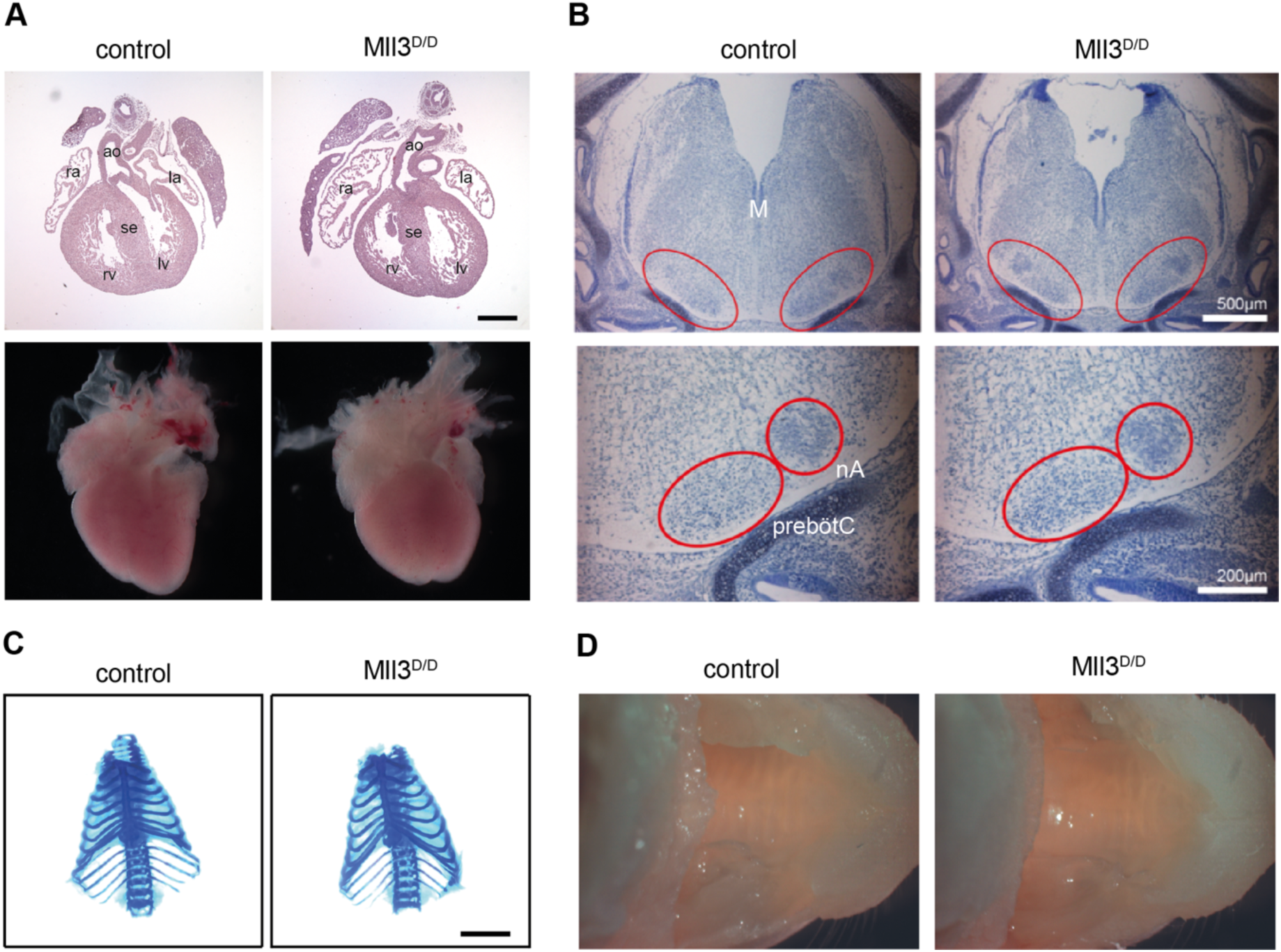
No morphological differences in Mll3 mutant fetuses. (A) Four chamber view of the fetal heart at E15.5 (top), morphology of the fetal heart at E18.5 (down). ra = right atrium, la = left atrium, rv = right ventricle, lv = left ventricle, se = intraventricular septum, ao = aorta. Scale bar 500 μm. (B) Overview of the brain stem at E15.5 (top) and nuclear enriched region of pre-Bötzinger complex and nucleus ambiguus (down) stained with Nissl. The size of these two regions was comparable among all genotypes. M = medulla, nA = nucleus ambiguus, preBötC = pre-Bötzinger complex. (C) Morphology of the rib cage after Alizarin red/Alcian blue staining at E18.5. Scale bar 5 mm. (D) Ventral view of the upper portion of the head showing the roof of the mouth at E18.5. The palate is intact in fetuses of different genotype.

**Fig. S5.**
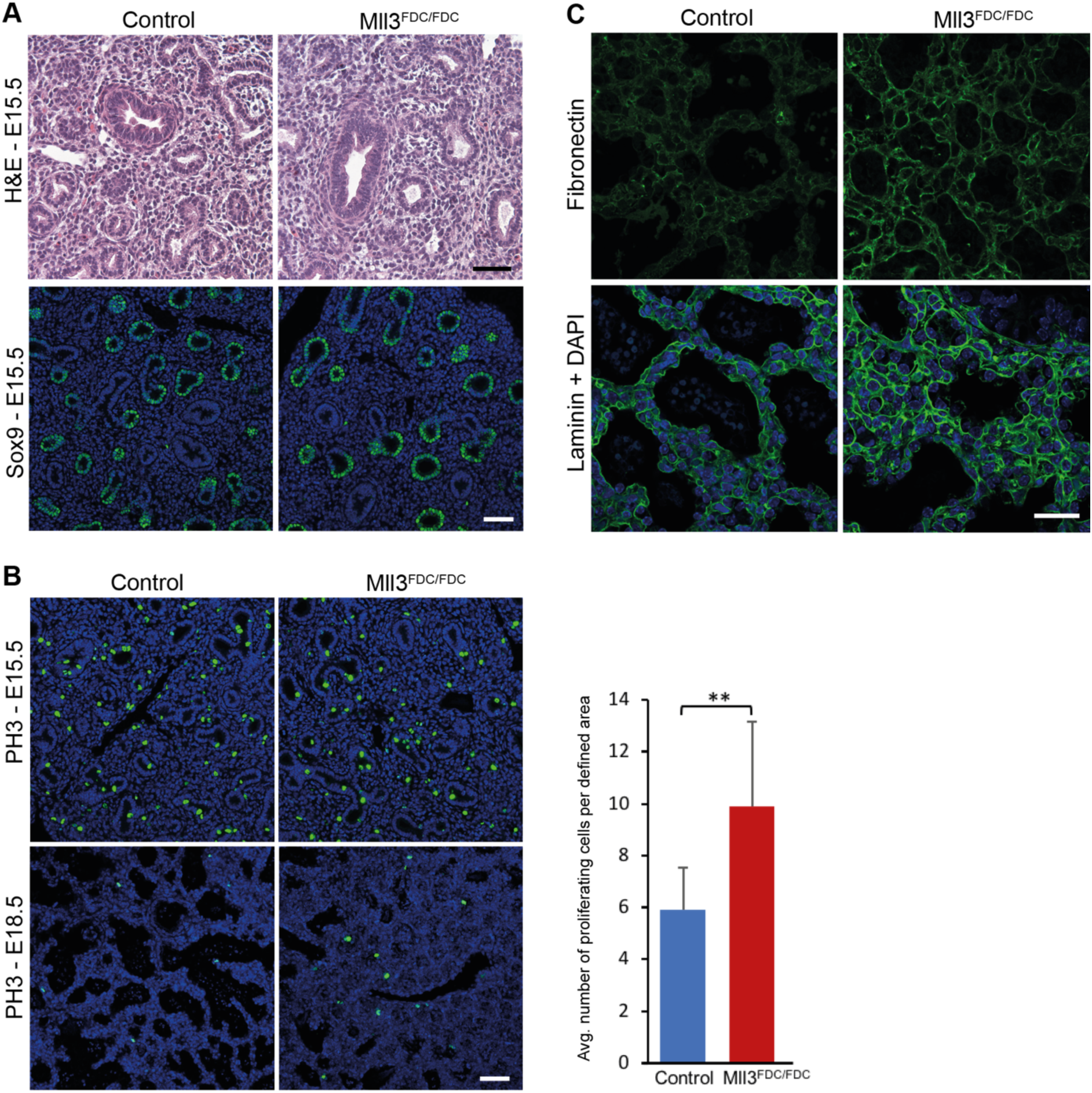
Increased proliferation and extensive deposition of extracellular matrix proteins in *Mll3^KO^* lungs. (A) H&E staining on sections of E15.5 lungs showed no morphological differences between control and *Mll3^KO^* littermates. Sox9 expression is seen in the distal epithelial tips of control and *Mll3^KO^* lungs. Scale bar 250 μm. (B) No difference in proliferation was observed between control and *Mll3^KO^* lungs of E15.5 fetuses after PH3 staining (top). At E18.5 an increase in proliferation is observed in *Mll3^KO^* lungs (bottom). Scale bar 250 μm. PH3 positive cells from E18.5 lung sections from control (n=3) and *Mll3^KO^* (n=3) lungs were counted. Four defined areas (each 1.8 mm^2^) were analysed. Mean ± s.d. is shown (**p <0.01 as calculated by unpaired t-test). (C) Extracellular matrix protein deposition of fibronectin and Lama1 in the basal lamina of *Mll3^KO^* lung is more compared to control littermate. Scale bar 25 μm.

**Fig. S6.**
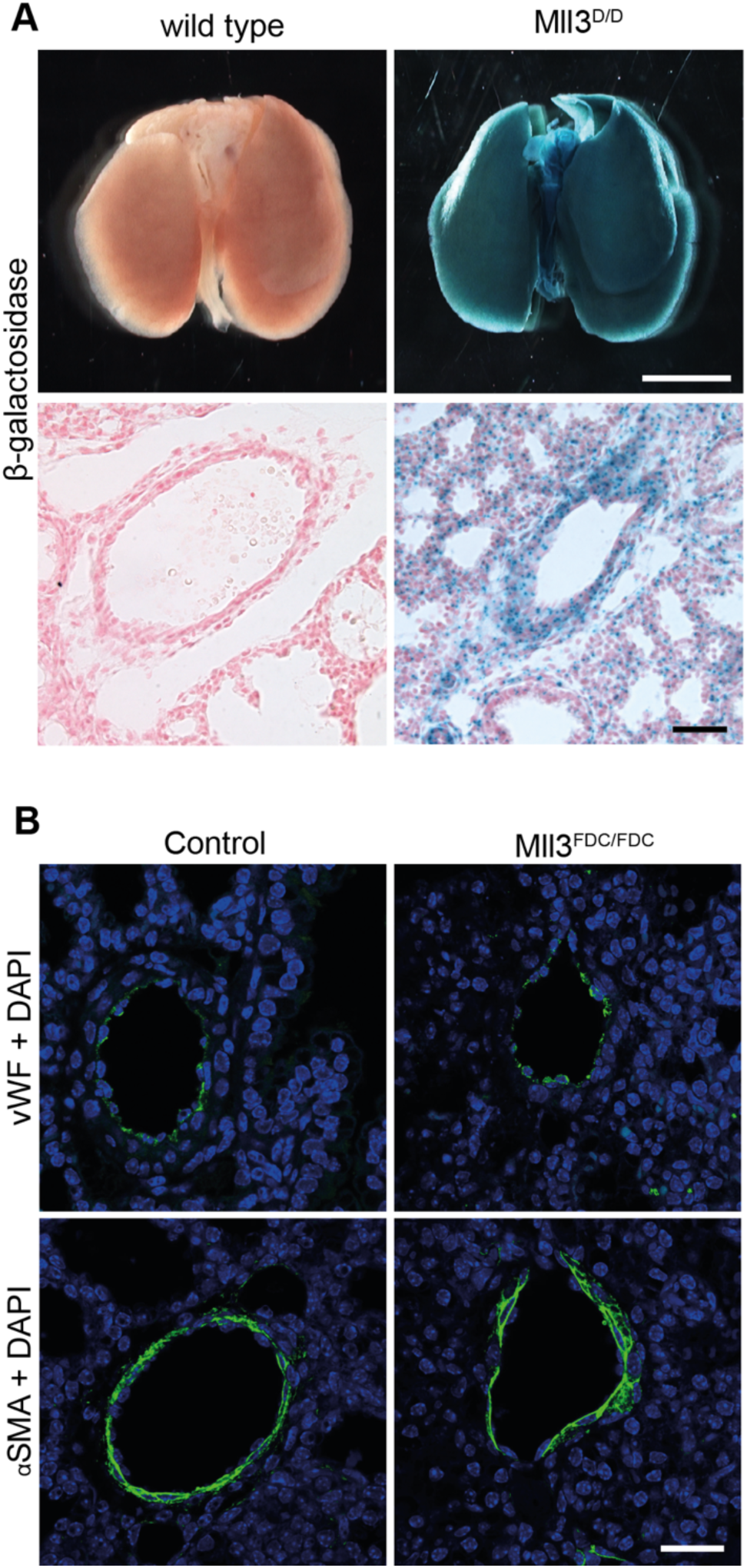
Normal lung vasculature at E18.5. (A) β-galactosidase staining reflecting MLL3 expression on whole lung (top) and lung sections (bottom). MLL3 expression in close proximity to pulmonary vessels demonstrated by β-galactosidase staining on wildtype and Mll3^D/D^ lung sections. Scale bar 500 μm (whole lung) and 250 μm (lung section). (B) Expression of vWF in endothelial cells of the large blood vessels in control and *Mll3^KO^* lungs. Smooth muscle surrounding the large blood vessels marked by expression of αSMA in control and *Mll3^KO^* lungs. Scale bar 25 μm.

**Fig. S7.**
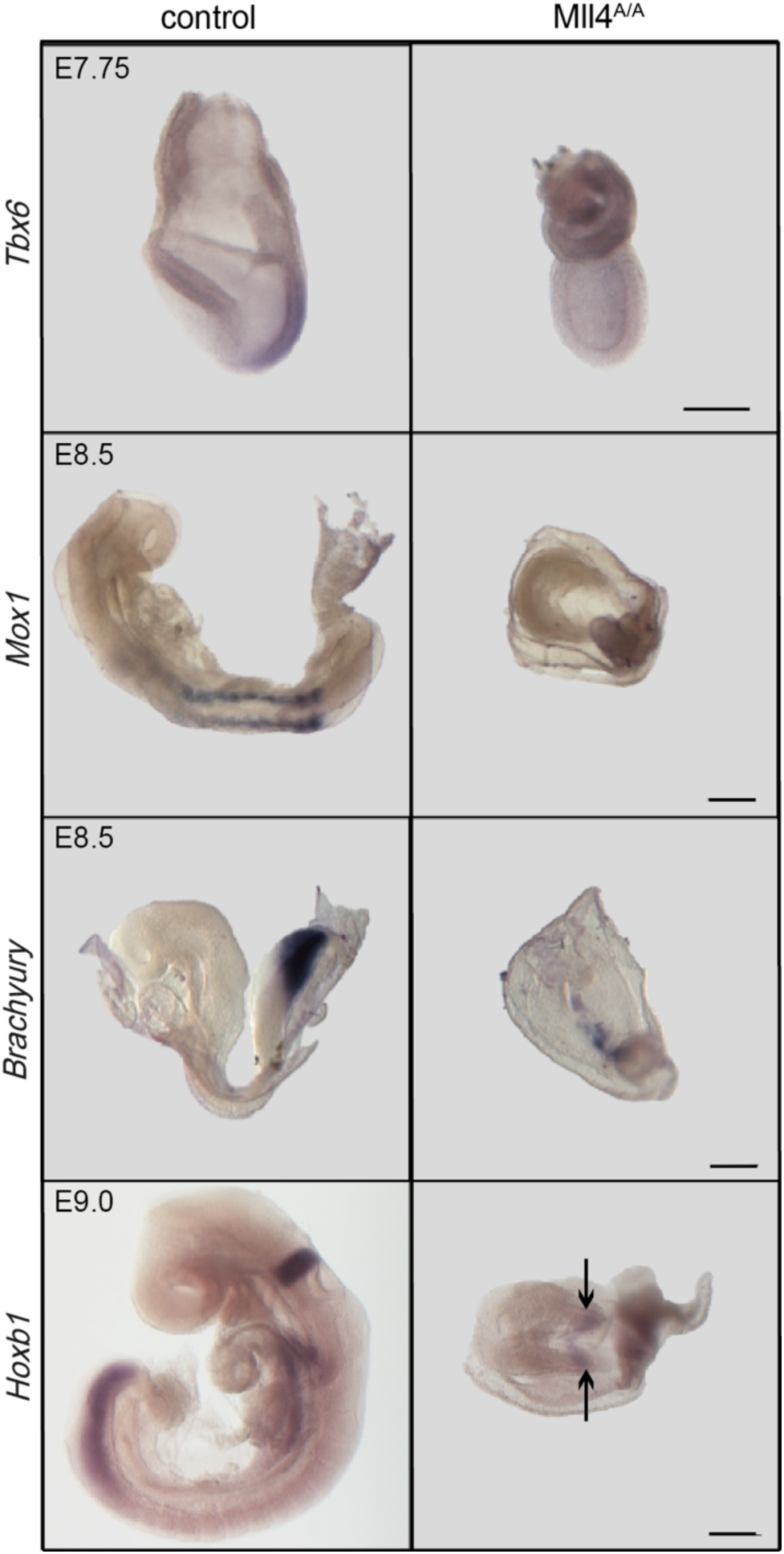
Defective gastrulation after loss of MLL4. Whole mount *in situ* hybridization of *Tbx6* (mesodermal wing marker) at E7.75, *Mox1* (somite marker) at E8.5, *Brachyury* at E8.5, and *Hoxb1* at E9.0 on embryos from *Mll4^A/+^* intercrosses. Arrows point towards the prospective rhombomere 4. Scale bars 250 μm.

**Table S1.**
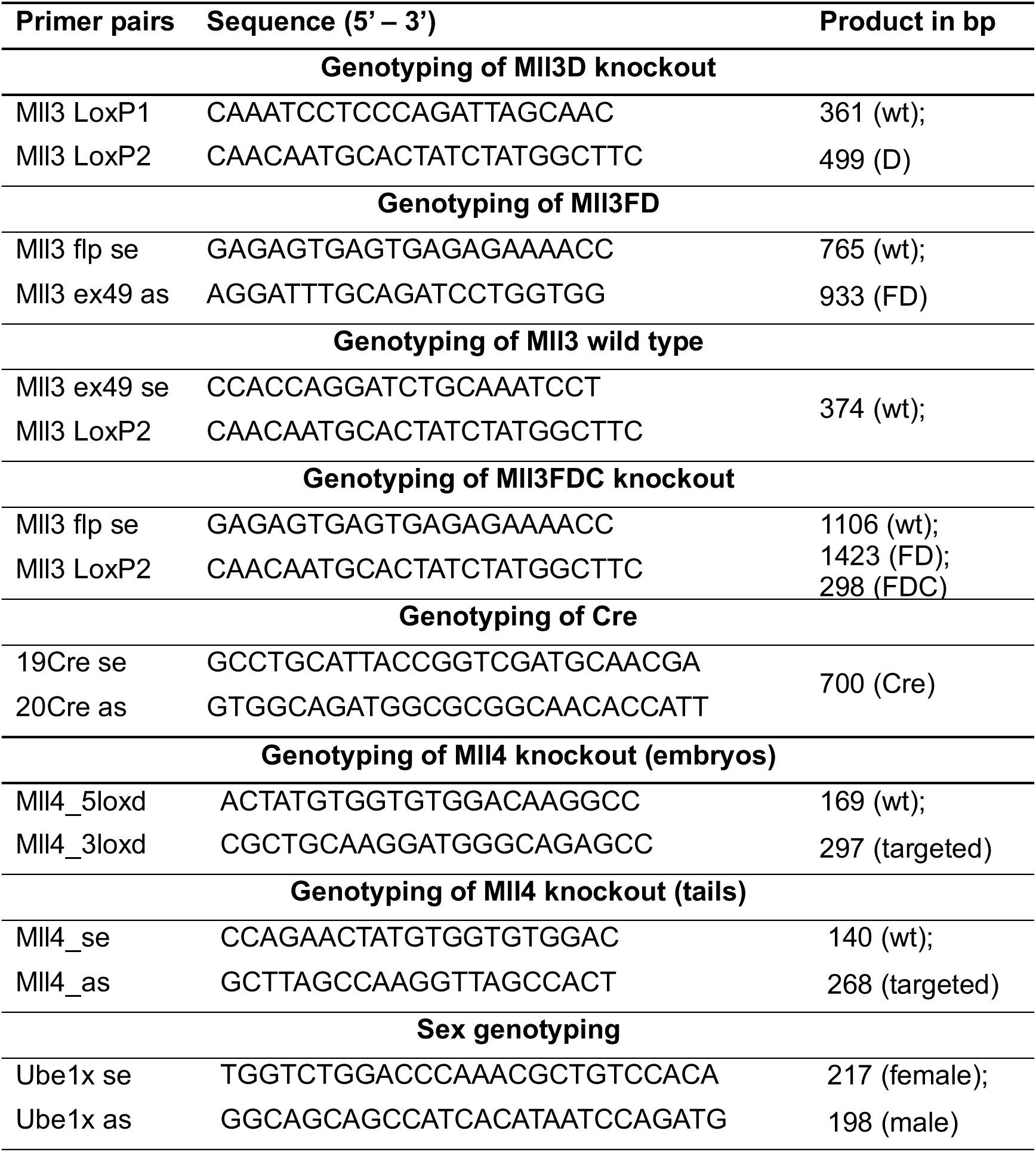
**Primers for genotyping**

**Table S2:**
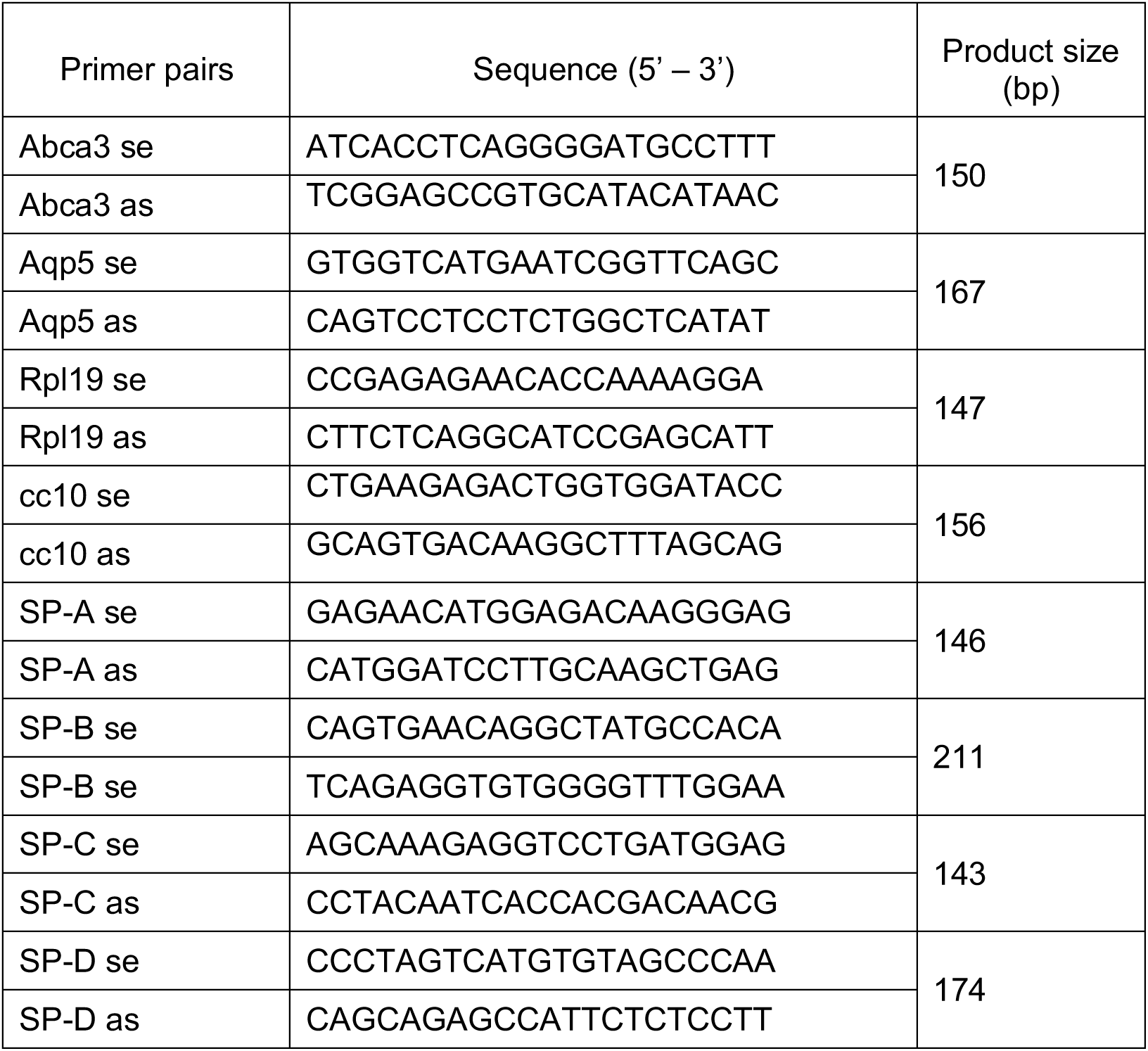
Primers for qRT-PCR

**Table S3:** Antibodies for Western blot (WB), immunofluorescence (IF) and immunohistochemical (IHC) staining

| **Antibodies** | **Company, Cat. Number** | **Dilution** |
| --- | --- | --- |
| **Primary antibodies** |  |  |
| **Rabbit anti-Hex** | Dr. Go Shioi | 1:2000 IF |
| Rabbit anti-cc10 | Santa Cruz, sc-25554 | 1:100 IF |
| Rabbit anti-MII4 | Diagenode, C15310100 | 1:500 WB |
| Rabbit anti-H3K4me1 | Diagenode, C15410037 | 1:2000 IHC |
| Rabbit anti-H3K4me2 | Diagenode, pAb-035-050 | 1:2500 IHC |
| Rabbit anti-H3K4me3 | Abcam, ab8580 | 1:1000 IHC |
| Rabbit anti-phospho-H3S10 | Upstate, 06-570 | 1:500 IF |
| Rabbit anti-Aquaporin 5 | Abcam, ab78486 | 1:800 IF |
| Rabbit anti-Fibronectin | Abcam, ab23750 | 1:500 IF |
| Rabbit anti-Lama1 | Sigma, L9393 | 1:100 IF |
| Rabbit anti-αSMA | Abcam, ab5694 | 1:100 IF |
| Rabbit anti-Lysozyme | Dako, A0099 | 1:500 IF |
| Rabbit anti-Sox9 | Millipore, AB5535 | 1:500 IF |
| Rabbit anti-von Willebrand Factor | Abcam, ab9378 | 1:100 IF |
| Mouse anti-ZO-1 | Thermo Scientific, 33-9100 | 1:500 IF |
| Rat anti-E-cadherin | Invitrogen, 13-1900 | 1:50 IF |
| **Secondary antibodies** |  |  |
| Biotinylated goat anti-Rabbit IgG | Vector, BA-1000 | 1:500 IHC |
| Goat anti-Rabbit IgG-HRP | Thermo Scientific, 31460 | 1:20000 WB |
| Goat anti-Rabbit IgG-CFL 488 | Santa Cruz, sc-362262 | 1:500 IF |
| Goat anti-mouse IgG Alexa Fluor 488 | Invitrogen, A11029 | 1:500 IF |
| Donkey anti-rat IgG Alexa Fluor 488 | Invitrogen, A21208 | 1:500 IF |

